# Mechanism of preferential complex formation by Apoptosis Signal-regulating Kinases

**DOI:** 10.1101/693663

**Authors:** Sarah J. Trevelyan, Jodi L. Brewster, Abigail E. Burgess, Jennifer M. Crowther, Antonia L. Cadell, Benjamin L. Parker, David R. Croucher, Renwick C.J. Dobson, James M. Murphy, Peter D. Mace

## Abstract

Apoptosis signal-regulating kinases (ASK1–3) are activators of the P38 and JNK MAP kinase pathways. ASK1–3 form oligomeric complexes known as ASK signalosomes that initiate signalling cascades in response to diverse stress stimuli. Here we demonstrate that oligomerization of ASK proteins is driven by previously uncharacterised sterile-alpha motif (SAM) domains that reside at the C-terminus of each ASK protein. SAM domains from ASK1–3 have distinct behaviours: ASK1 forms unstable oligomers, ASK2 is predominantly monomeric, and the ASK3 SAM domain forms a stable oligomer even at low concentration. In contrast to their isolated behaviour, the ASK1 and ASK2 SAM domains preferentially form a stable heterocomplex. The crystal structure of the ASK3 SAM domain, small-angle X-ray scattering, and mutagenesis suggests that ASK3 oligomers and ASK1-ASK2 complexes form discrete quasi-helical rings, via the mid-loop–end-helix interface. Preferential ASK1-ASK2 binding is consistent with mass spectrometry showing that full-length ASK1 forms heterooligomeric complexes incorporating high levels of ASK2. Accordingly, disruption of SAM domain-association impairs ASK activity in the context of electrophilic stress induced by 4-hydroxy-2-nonenal. These findings provide a structural template for how ASK proteins assemble foci to drive inflammatory signalling, and reinforce that strategies targeting ASK kinases should consider the concerted actions of multiple ASK family members.

## Introduction

Mitogen-activated protein kinase (MAPK) cascades are ubiquitous in eukaryotes as a means of sensing and responding to stressors. In humans, the JNK and P38 MAP kinases are activated by upstream MAP kinase kinases (MAP2Ks), which are in-turn activated by a diverse group of MAP3Ks. Although the activation of both MAPKs and MAP2Ks by phosphorylation is well understood, MAP3Ks are less well characterised. This imbalance likely stems from the fact that while MAPKs and MAP2Ks are activated by relatively well-defined upstream kinases, stress-activated MAP3Ks must recognise and respond to a wide range of stressors so have more diverse regulation that remains to be characterised at the molecular level.

Apoptosis signal-regulated kinases (ASKs) are a group of MAP3Ks that respond to a broad variety of chemical, physical and inflammatory stimuli. In humans there are three ASK-family kinases (ASK1–3; MAP3K5, MAP3K6 and MAP3K15, respectively). ASK1 has been intensively studied following the initial discovery of its activation in response to TNFα, promoting cell death (*1*). Subsequently, roles for all three ASK kinases have been defined in various biological pathways and disease states. For instance, ASK1 is now well established in the response to oxidative stress and inflammatory cytokines (*2, 3*). ASK1 and ASK2 are required for effective responses to viral infection (*4–6*), to prime NLR inflammasomes following challenge by bacterial infection (*7*), and together act to mediate neutrophillic dermatitis (*8*). In simplified terms, it appears that ASK1 and ASK2 in isolation are each able to promote some level of P38/JNK activation and stress response, but their concerted action generates a broader inflammatory response, and cell death. ASK3 apparently has a more specialised role in sensing and responding to osmotic pressure and regulation of blood pressure, specifically in the kidney via the WNK1 pathway (*9*).

Recently, ASK1 has generated significant interest due to the relevance of ASK1 to disease and the availability of specific inhibitors, in particular Selonsertib (*10*). Activating mutations of ASK1 occur in melanoma (*11*), and inhibition of ASK1 has shown benefit in gastric cancers (*12, 13*). Most notably, ASK1 is a relevant target in non-alcoholic steatohepatitis (NASH; (*14*)). ASK1 inhibition with Selonsertib has shown promising results up to phase two clinical trials (*10*), and inhibitors derived from Selonsertib also reduce fibrosis caused by kidney inflammation (*15*). Despite their clinical relevance, a structural understanding of ASK protein complexes beyond the well-conserved catalytic kinase domain is limited. The ASK1 kinase domain structure was first solved in 2007 (*16*), and subsequently crystal structures of small-molecule inhibitors in complex with the kinase have become available. However, ASK1–3 are each greater than 1300 amino acids in length and the precise mechanisms linking their conserved architecture—where the central kinase domain is flanked by large N- and C-terminal regulatory domains (Figure 1A)—to kinase activity remain unclear.

**Figure 1:**
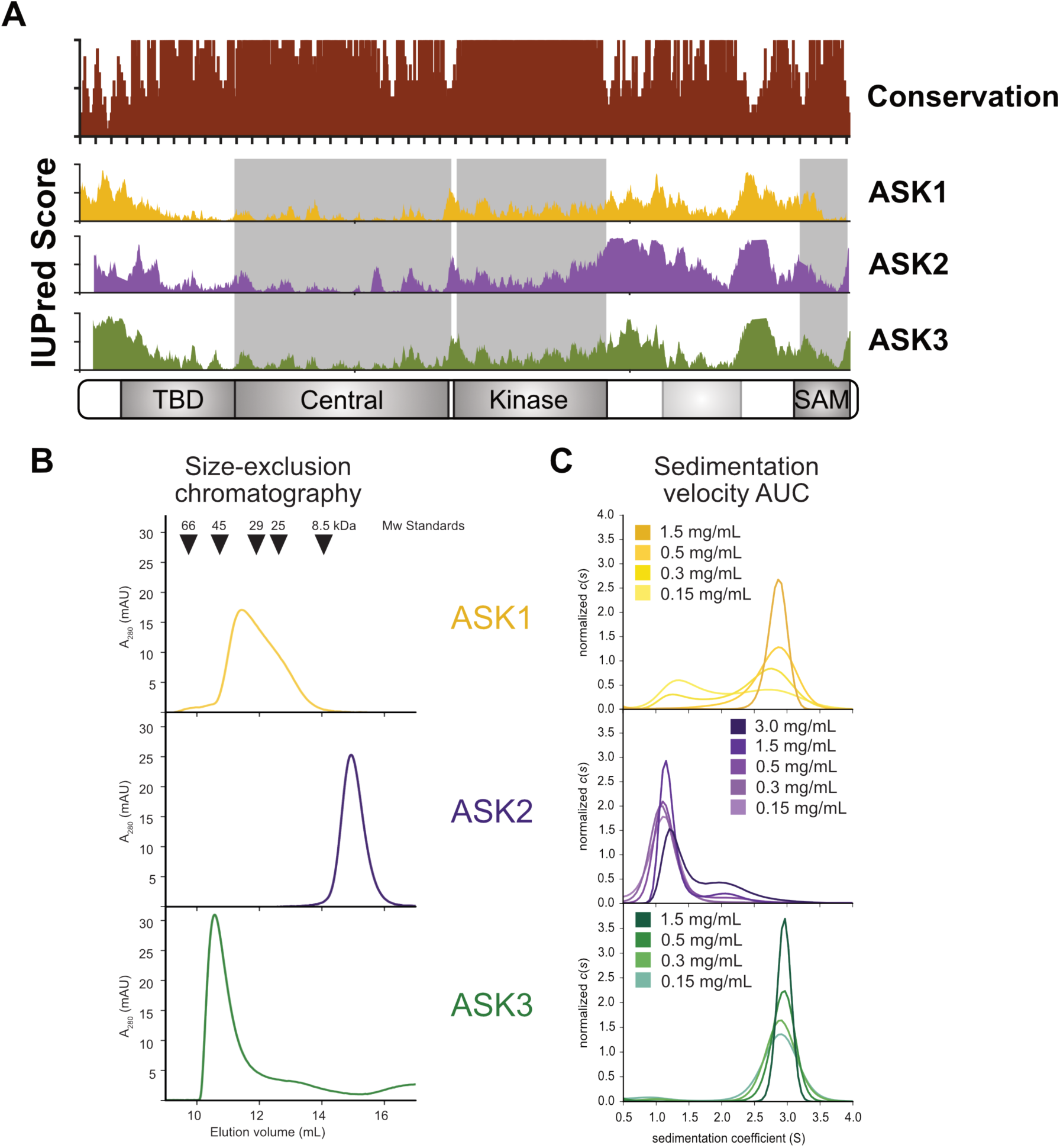
ASK1, ASK2 and ASK3 C-terminal domains have different oligomerisation propensity. (A) Overview of domain architecture, and conservation of ASK1–3. (B) Size-exclusion chromatography of ASK1(1290–1374), ASK2(1216–1288) and ASK3(1241–1313), corresponding to regions of respective ASK proteins labelled ‘SAM’ domain in panel A. (C) Sedimentation velocity AUC analysis of ASK1, 2 and 3 domains at indicated concentrations spanning between 0.15 and 3.0 mg/mL (15–365 µM).

The current model of ASK1 regulation invokes constitutive oligomerisation mediated through the C-terminal region, in parallel with stimuli-dependent regulation of ASK signalling via the N-terminus (*17*). Many of the signalling molecules that are proposed to regulate stress-induced activation of ASK1 interact via its N-terminal domains. There are further outstanding questions regarding the interaction of regulatory and oligomerising domains of ASK proteins. It is not clear if regulation of substrate recruitment and priming, through a domain just N-terminal to the kinase (*18*), occur in an intra- or inter-molecular manner. Likewise it is not known if dimers reported for the isolated kinase domain of ASK1 impact kinase function in the context of full-length protein (*16*). Moreover, the C-terminal region of ASK proteins is clearly important for signalosome formation and activity, but the structural mechanism of assembly and how this relates to oligomerisation of different ASK-type kinases remains to be determined.

Here we present the crystal structure of the C-terminal domain of ASK3, which adopts a sterile alpha motif—a fold classically shown to mediate protein-protein interactions that had not previously been described in ASK proteins. Interrogating the behaviour of C-terminal domains of ASK1–3 using a variety of methods in solution and full length ASK1/2 in cells uncovers distinct behaviours of the C-terminal domains from the three ASK proteins, which impact protein complex assembly and activity. This data provides a structural basis for previous observations regarding ASK protein oligomerisation, and functional cooperativity of ASK proteins in a variety of biological settings.

## Results

### ASK1–3 C-terminal domains have divergent oligomerisation propensities

Sequences C-terminal to the ASK1 kinase domain are known to play roles in binding regulatory proteins (*19, 20*), and facilitating interactions between ASK proteins to generate oligomeric ASK signalosomes (*17*). While a coiled-coil region is predicted near the C-terminus of each ASK protein, the precise structural architecture of the C-terminal portion of ASK proteins is unclear. To gain insight into the mechanism of oligomerisation, we expressed C-terminal fragments from ASK1, ASK2, and ASK3. Regions of ASK1(1039– 1374), ASK1(1237–1374), ASK2(988–1288), and ASK2(1156–1288) that incorporate the predicted C-terminal coiled-coil (residues 1245–1285 in ASK1) were all highly insoluble when expressed alone or co-expressed, in either *E. coli* or Sf9 insect cells. In contrast, shorter constructs comprising ASK1(1290–1374), ASK2(1216–1288) and ASK3(1241– 1313) (9.8, 8.2, 8.5 kDa, respectively) all readily expressed in a soluble form in *E. coli*.

Assessment of the soluble C-terminal portions of ASK proteins using analytical size-exclusion chromatography (SEC) and analytical ultracentrifugation (AUC) revealed that these smaller fragments themselves had the ability to form oligomers. However, each exhibited a distinct behaviour. SEC showed an unstable ASK1 oligomer that existed in an equilibrium between multiple oligomeric states, even at concentrations as high as 200 µM (Figure 1B). Sedimentation velocity AUC corroborated this result, whereby between 15–150 µM ASK1 formed concentration-dependent oligomers of two distinct sizes (Figure 1C; Supp. Table 1). In comparison, ASK2 was a single species on SEC, eluting with an apparent mass consistent with a monomer (Figure 1B). AUC also showed that the ASK2 exists almost exclusively as a monomer, only exhibiting a minor dimer species when analysed at a concentration of 365 µM (3 mg/mL). Finally, ASK3 forms a large and stable oligomer, with an apparent mass of ∼54 kDa assessed from molecular weight standards (Figure 1B). Analysis of ASK3 using AUC suggested a single oligomeric state over a 10-fold concentration range (Figure 1C; Supp. Table 1), allowing for a good mass estimation from the sedimentation velocity experiment. When measured between 0.15 and 1.5 mg/mL, the calculated molecular weight values fell in a range between 41.3 and 47 kDa, generally between the mass of an ASK3 pentamer or hexamer, which would have a theoretical mass of ∼42.5 kDa or 51 kDa respectively.

**Table 1:** Crystallographic data

|  |  |  |
| --- | --- | --- |
| Data Collection |  |  |
| ASK3 SAM domain | Native | Iodide |
| Beamline | AS-MX2 | AS-MX2 |
| Wavelength (Å) | 0.9357 | 1.456 |
| Resolution (Å) | 48.13–1.80 (1.83–1.80) | 48.11-2.62 (2.74-2.62) |
| Space group | $P2_12_12_1$ | $P2_12_12_1$ |
| Unit cell parameters<br>a, b, c (Å)<br>$\alpha$, $\beta$, $\gamma$ (°) | 45.42, 53.56, 96.26<br>90, 90, 90 | |
| R<sub>merge</sub> (outer shell) | 0.053 (0.985) | 0.114 (0.474) |
| R<sub>pim</sub> (outer shell) | 0.044 (0.787) | 0.059 (0.365) |
| Mean I/σI (outer shell) | 11.5 (1.1) | 11.2 (2.3) |
| Completeness (outer shell) | 99.4 (98.3) | 99.0 (93.2) |
| Multiplicity (outer shell) | 4.2 (4.4) | 7.5 (4.2) |
| Total No. of reflections | 94216 | 56242 |
| No. of unique reflections | 22425 | 7518 |
| CC(1/2) (outer shell) | 0.998 (0.455) | 0.997 (0.749) |
| Wilson B factor (Å<sup>2</sup>) | 35.05 |  |
| Refinement Statistics |  |  |
| R<sub>cryst</sub> | 0.2130 |  |
| R<sub>free</sub> | 0.2614 |  |
| rmsd for bonds (Å) | 0.013 |  |
| rmsd for angles (deg) | 1.30 |  |
| rmsd for chiral volume (Å<sup>3</sup>) | 0.072 |  |
| No. of protein atoms | 1688 |  |
| No. of solvent atoms | 122 |  |
| Average B-factor overall (Å<sup>2</sup>) | 39.36 |  |
| Average main chain B-factor (Å<sup>2</sup>) | 38.93 |  |
| Average side chain and solvent B-factor (Å<sup>2</sup>) | 45.27 |  |
| Ramachandran plot statistics (%) |  |  |
| Favoured regions | 99.5 |  |
| Allowed regions | 0 |  |
| Outliers | 0.5 |  |

To further interrogate the soluble C-terminal domains of ASK1–3 we analysed their primary sequences using sequence profile matching (*21*), which suggested significant homology to sterile-alpha motif (SAM) domains from various proteins—including P63, Tankyrase, and yeast MAPK-related proteins Ste11 and Ste50 (*22–25*). While there has been some reference to a predicted SAM domain in ASK1 (*26*), the SAM designation does not appear in major fold-prediction databases, and the same region has also been noted to contain a ubiquitin-like sequence motif (*26, 27*). Because SAM domains are versatile interaction modules that mediate both protein-protein and protein-DNA interactions, a C-terminal SAM domain would make an ideal candidate to mediate ASK oligomer formation.

### The ASK3 C-terminal domain is a Sterile-Alpha Motif (SAM) domain

To gain further insight into the oligomerisation mechanism of the ASK1–3 C-terminal domains we pursued structural studies. Crystallisation trials of the soluble C-terminal domains from the three ASK proteins yielded crystals of ASK3(1241–1313), from which the structure was solved using a combination of SIRAS and MR-SAD (see methods; Figure 2A). The structure was refined against native diffraction data to a resolution of 1.8 Å, and has excellent geometric parameters (Table 1). There are three molecules of ASK3(1241–1313) in the asymmetric unit; with the ASK3 polypeptide chain defined from residues 1241–1308 in two molecules, and 1241–1305 in the remaining molecule.

**Figure 2:**
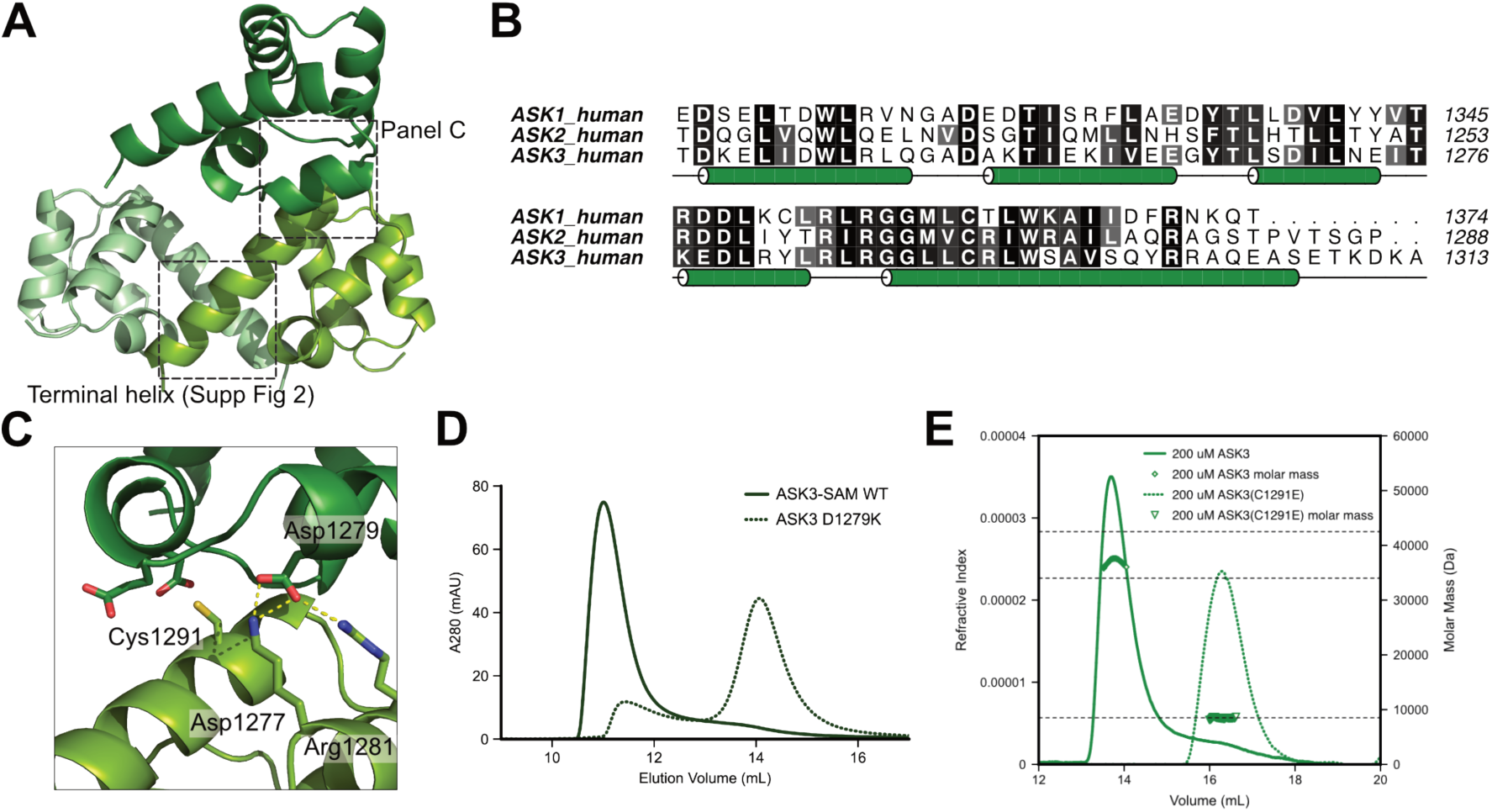
Structure of the ASK3 SAM domain. **(A)** Cartoon representation of the crystal structure of ASK3(1290-1374) displaying the three monomers within the asymmetric unit. The ML-EH interface is indicated with a dashed box. **(B)** Alignment of SAM domains of human ASK1–3. **(C)** Close-up view of wild-type residues within the dashed areas of the ML-EH interface. **(D)** Size-exclusion chromatography trace indicating the disruption of the wild-type ASK3-SAM oligomer as a result of a aspartate to lysine mutation at position 1279 (ASK3(D1279K)). **(E)** SEC-MALS data measuring the molar mass of a cysteine to glutamate mutant at position 1291 (ASK3(C1291E)), relative to the wild-type oligomer.

Consistent with bioinformatic prediction, the C-terminal domain of ASK3 adopts the classical five-helix fold of the sterile-alpha motif domain (SAM). The sequence of ASK1 and ASK2 are 53 and 37 % identical in the equivalent regions to the solved structure of ASK3 (Figure 2B). The highest levels of conservation are concentrated in hydrophobic core residues, and both ASK1 and ASK2 are also predicted to contain five helices. Therefore, we propose that the three ASK proteins all possess a similar SAM-fold at their C-terminus. Although we grew crystals of the SAM domain of ASK1, the crystals did not diffract sufficiently for structure determination.

### Oligomer formation by the ASK3 SAM domain

With three ASK3 molecules in the asymmetric unit, there are several interfaces through which ASK oligomers may form. Based on the crystal contacts, we observed three possibilities (Supp. Fig. 1): the mid-loop:end-helix (ML-EH) interaction that has been observed for SAM domains from diverse protein families (Figure 2A/C); a symmetrical interaction formed by the C-terminal helix of ASK3, and a symmetrical interaction though the surface of α1 and α2 with a neighbouring asymmetric unit. We generated a suite of mutant ASK3 SAM domains to deduce which of the interfaces observed in the crystal lattice corresponds to the oligomerisation interface in solution. Examining mutants by size-exclusion chromatography, we identified the ML-EH interaction as the crucial site for oligomerisation. Strikingly, the (ML-EH) mutant D1279Q eluted with an apparent mass of 13 kDa, close to that of a monomer (Figure 2C/D). Mutation of residues at the C-terminal helix led to ambiguous results: ASK3 Y1300Q caused a shift towards smaller apparent mass of 22 kDa, and Q1304A did not affect the wild-type ASK3 SAM oligomer (Supp. Fig. 2). The V1262N and K1242E mutants at the α1/α2 interface did not disrupt the oligomer (Supp. Fig. 2), indicative of purely crystallographic contacts through α1/α2. Thus, we cannot completely discount the role of the C-terminal-helix interaction, but the ML-EH interaction is indispensible for ASK3 oligomer formation and a single point mutant at this interface most effectively disrupts the ASK3 complex. While there is some possibility of a C-terminal interaction ASK3, we can effectively exclude such an interaction for ASK1 based on several criteria: an equivalent ASK1 mutation (F1369Q) behaves in an identical manner to WT protein (Supp. Fig. 3A); ASK1 has a truncated C-terminal helix relative to ASK3 that would disrupt half of the interacting surface (Supp. Fig. 3B); and two ASK3 residues from α3 that pack against the C-terminal helix (Ser1269 and Asn1273) are not conserved in ASK1 (Supp. Fig. 3B).

**Figure 3:**
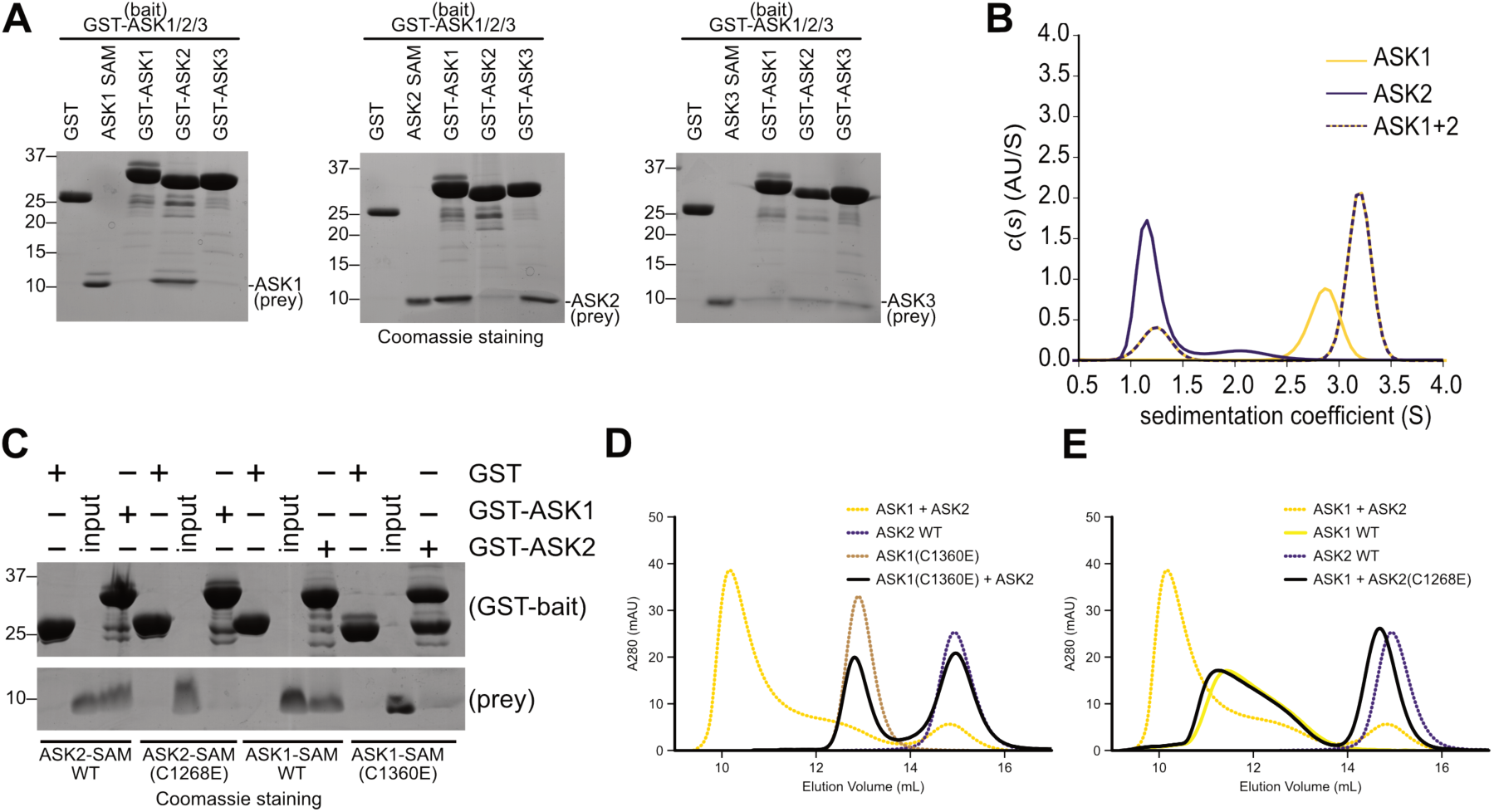
Heterotypic interactions between ASK SAM domains **(A)** GST-pulldown experiments measuring the ability of GST-ASK1, -ASK2, and -ASK3 SAM domains to pull down untagged SAM domains from ASK1, ASK2 and ASK3 respectively **(B)** Sedimentation velocity analytical ultracentrifugation of the isolated SAM domains of ASK1, ASK2 (1.5 mg/mL) and an equimolar mixture of the two (at 1.5 mg/mL each). **(C)** GST-pulldown experiments measuring the ability of WT GST-ASK1 and GST-ASK2 SAM domains to pull down either WT, or Cysteine mutant ASK1 and ASK2 SAM domains. **(D)** Analytical size-exclusion chromatography comparing the ability of WT and C1360E ASK1 SAM domain to form a higher-order oligomer with the WT ASK2 SAM domain. **(E)** Analytical size-exclusion chromatography comparing the ability of WT and C1268E ASK2 SAM domain to form a higher-order oligomer with the WT ASK1 SAM domain.

To further probe the role of the ML-EH interface in ASK interactions, we created an additional mutation on the opposite side of the interface in both ASK3(C1291E) and ASK1(C1360E), and used SEC-MALS to test their ability to oligomerise. Consistent with an indispensable role for the ML-EH interface, ASK3(C1291E) had a calculated mass of 8.5 kDa (Figure 2E), and ASK1(C1360E) had a calculated mass of 13.4 kDa (Supp. Fig. 3C). The wild-type ASK2 SAM domain had a calculated mass of 8.6 kDa in SEC-MALS, even though it eluted in a notably different position relative to ASK1/ASK3 monomers (Supp. Fig. 3). This may indicate that ASK2 has more compact monomeric hydrodynamic radius and/or different flexibility at the C-terminal tail relative to ASK1 and ASK3.

From these data, we conclude that weak oligomerisation of the ASK1 SAM domain and stable oligomer formation by the ASK3 SAM domain requires the ML-EH interface, because single point mutants that disrupt the ML-EH behave as a monomeric proteins.

### ASK1-ASK2 form heterotypic complexes via the ML-EH interface

ASK1 and ASK2 have previously been reported to associate through their C-terminal domains (*4*), and endogenous ASK1 and ASK2 have been reported to exist in complex with one another at an equal ratio (*28*). We next sought to test whether such heterotypic association could be driven through the isolated SAM domains of each ASK protein. As an initial measure, we prepared GST-fused forms of the SAM domain from each ASK protein, and tested the ability of each to pulldown the other respective SAM domains (Figure 3A). Strikingly, we observed that ASK1 SAM domain clearly only associated with the ASK2 domain, but not untagged ASK1, or ASK3 SAM. The ASK2 SAM domain was not able to pull down its own untagged form—consistent with its monomeric behaviour in solution (Figure 1B/C; Supp. Table 1)—but could readily pull down ASK1 and ASK3 (Figure 3A). ASK3 on the hand showed only weak interactions with untagged SAM domains from any ASK protein (Figure 3A). The scarcity of interactions by ASK3 could be because ASK3 readily forms oligomers over a range of concentrations (Figure 1C), and thus GST-ASK3 SAM is unable to incorporate further untagged ASK3 SAM.

To further characterise the heterotypic association of ASK1-ASK2 we used sedimentation velocity AUC (Figure 3B; Supp. Table 1). Consistent with GST-pulldowns, the ASK1-ASK2 mixture readily associated, and formed a defined oligomer. Such behaviour is in marked contrast to the weak homotypic association of ASK1 and ASK2 SAM domains, and suggests that the two domains from ASK1 and ASK2 preferentially oligomerize into a larger complex. This behaviour suggests that the SAM domains of each protein contribute markedly to the heterotypic ASK1-ASK2 complexes previously reported (*4*–*8, 28*).

To ascertain if ASK1-ASK2 hetero-oligomers also use the ML_-_EH interface, we first tested the role of the conserved cysteine residue, previously shown to be essential for ASK3 oligomerisation and for weak ASK1 oligomerisation. GST-pulldown experiments clearly showed that mutating either Cys1360 of ASK1, or Cys1268 of ASK2 to glutamate ablated binding to the WT form of its partner protein (Figure 3C). To gain further information on the oligomerisation status of these mutant proteins we performed analytical-SEC (Figure 3D/E). We observed that ASK1(C1360E) showed no higher order complex formation when combined with WT ASK2 SAM domain, having precisely the equivalent elution time as when it was analysed by itself. The corresponding mutation in ASK2(C1268E) reinforced this observation, with the mixture of ASK1–ASK2(C1268E) SAM domains barely distinguishable from their isolated elution positions, showing no sign of higher-order complex assembly. Together, these interaction studies show that the SAM domains from ASK1 and ASK2 exhibit a preference to form heterotypic, rather than homotypic, higher-order oligomers through the ML-EH interfaces of each protein.

### SAM-mediated oligomers regulate activity and stoichiometry in cells

Having established that disrupting the ML-EH interface of either ASK1 and ASK2 abrogates SAM domain heterocomplexes, we sought to test the effects on activity. We transfected wild-type full-length, and disruptive mutations ASK1(C1360E) and ASK2(C1268E), into HEK293FT cells and challenged cells with the prototypic electrophilic stress 4-hydroxy-2-nonenal (HNE). As expected, HNE treatment of cells overexpressing wild-type full-length ASK1 induced significant phosphorylation of ASK1, indicating activation of kinase activity. The oligomer disrupted mutant ASK1(C1360E) showed decreased relative phosphorylation upon HNE stimulation, characteristic of impaired activity (Figure 4A/B). When transfected together, wild-type ASK1 and ASK2 show significantly lower basal levels of kinase activity, which is nonetheless activated by HNE. Based on the complete lack of signal in cells transfected with ASK2 alone (WT or C1268E; Figure 4A/B central lanes), the phospho-ASK signal is specific to ASK1, rather than cross-reactivity of the ASK1-phosphoT845 antibody. A plausible explanation for this observation is ASK1 in complex with ASK2 favours transphosphorylation (*4*), and therefore limits ASK1 autophosphorylation. Strikingly, transfection of ASK1(C1360E) and ASK2(C1268E) together leads to complete abrogation of basal phosphorylation, which is not stimulated by HNE treatment (Figure 4A/B). From these experiments, we conclude that oligomerisation of ASK proteins through their SAM domains is a core component of active signaling complex formation, which can be disrupted by single point mutations at their ML-EH interfaces.

**Figure 4:**
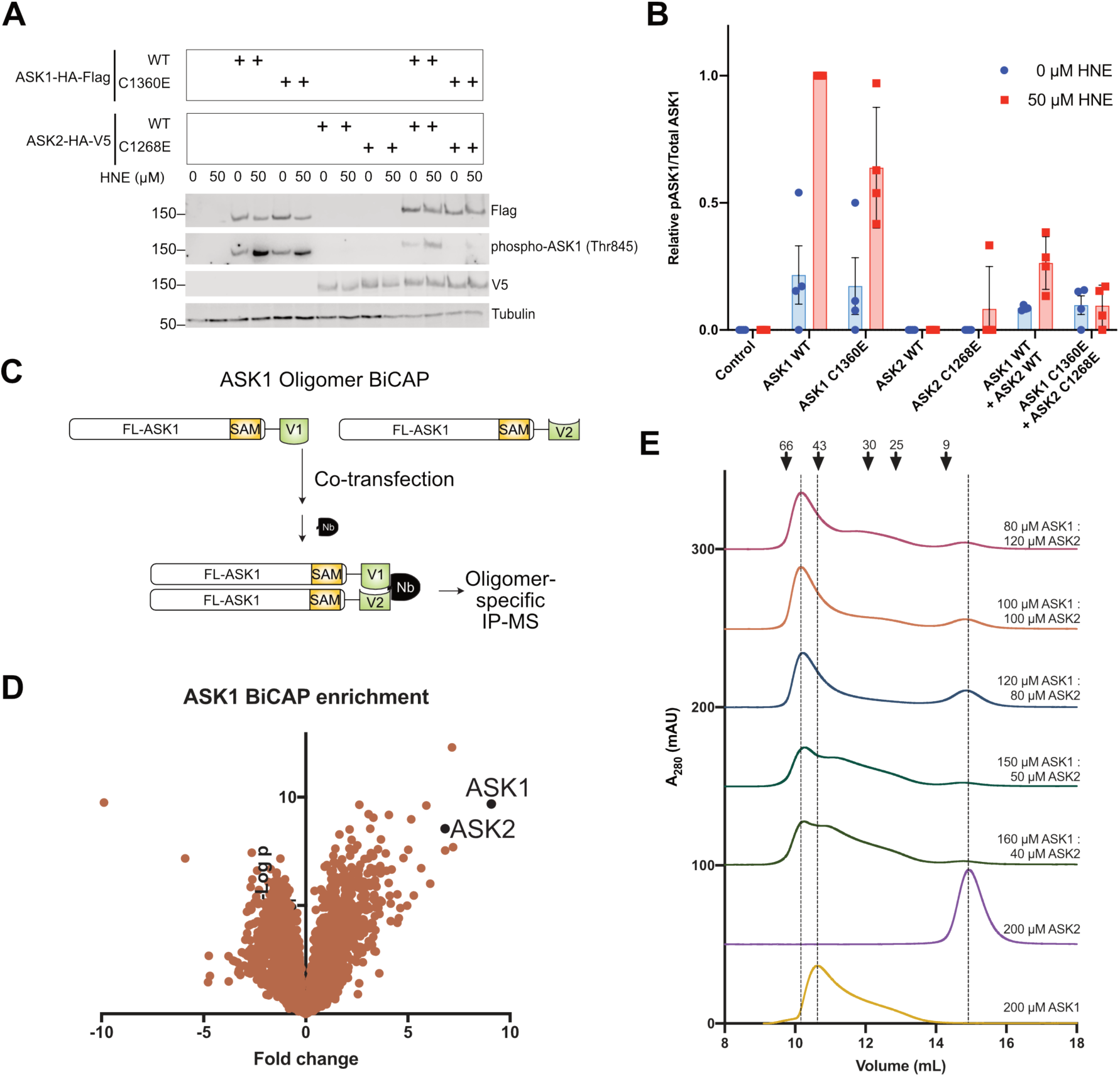
Role of the ML-EH interface in ASK signaling and stoichiometry **(A)** Analysis of HEK293 cells transfected with indicated constructs, either unchallenged or challenged with HNE (50 µM) for 1 hour. **(B)** Quantitation of the ratio between total ASK1 (assessed via Flag immunoblotting in Panel A) and phospho-Thr845 ASK1, four independent biological replicates. **(C)** Schematic illustration of the BiCAP system as applied to ASK1. **(D)** Waterfall plot of BiCAP MS/MS data following anti-GFP nanobody immunoprecipitation. Data is expressed as the fold-change over abundance calculated from a GFP only-transfected cell line treated in an equivalent manner control. **(E)** Analytical size-exclusion chromatography of mixtures containing indicated concentrations of the ASK1 and ASK2 SAM domains.

Elegant endogenous mass spectrometry studies have previously shown that ASK1 associates with near-stoichiometric amounts of ASK2 (*28*). We employed an orthogonal approach—bimolecular complementation affinity purification (BiCAP (*29, 30*))—to determine if stoichiometric association of ASK1-ASK2 occurs as part of larger hetero-oligomeric complex. For this system, we created two constructs of full-length ASK1 fused to the N-terminal (V1), and C-terminal (V2), portions of Venus fluorescent protein employed in BiCAP. As such, complexes immunoprecipitated using an anti-GFP nanobody must contain *at least* two molecules of full-length ASK1 (Figure 4C), rather than associating with any monomeric form of the protein, as may occur with a conventional immunoprecipitation. Partner proteins identified with multimeric ASK1 were identified by mass spectrometry. Remarkably, ASK2 was identified at an abundance of approximately 75% of ASK1 itself (Figure 4D; Supp. Table 2), even though ASK1 was overexpressed whilst ASK2 was expressed at endogenous levels. Such a result strongly suggests a selective incorporation of near-equal ratios of ASK1 and ASK2 into higher order ASK complexes. Also of note, ASK3 was also enriched in BiCAP analysis, as were several members of the ubiquitin ligase machinery (FbxW11, UBE2N/Ubc13). ASK proteins have previously been shown to undergo regulatory ubiquitination (*20, 31, 32*).

**Table 2:**
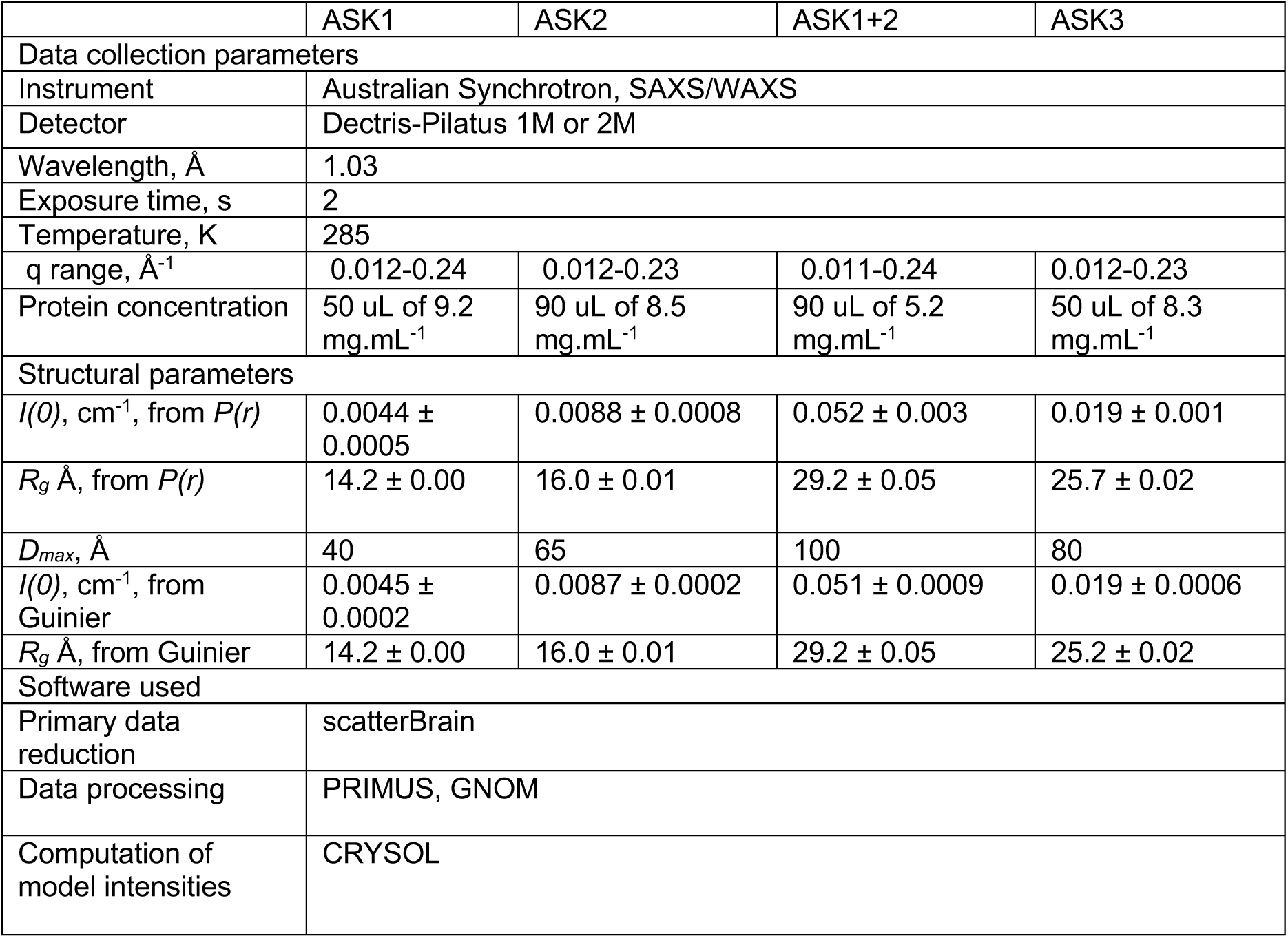
SAXS Data Parameters

Finally, to ascertain whether near equal ASK1-ASK2 stoichiometry observed in cells is recapitulated by isolated SAM domains we completed a series of analytical size-exclusion chromatography experiments with different ratios of ASK1 and ASK2 SAM domains. In these experiments, near equal ratios of ASK1 and ASK2 SAM domains (120:80, 100:100 µM ASK1:ASK2) led to the most homogenous higher order complexes, from a range ratios tested (Figure 4E; 200:0, 160:40, 150:50, 120:80, 100:100, 80:120, 0:200 µM, ASK1:ASK2, respectively). Together, these results suggest that SAM domains are a major determinant of ASK oligomeric state, promoting higher order complexes with near equal ratios of ASK1 to ASK2, that can be disrupted to decrease kinase activity in cells.

### Comparison of ML-EH surfaces in ASK paralogs

The ML-EH interaction occurs in several SAM domain complexes, including both discrete heterodimeric interactions and polymeric arrays of SAM domains. To name contrasting examples: CNK:HYP, and EPH:SHIP2 SAM domains that form a heterodimer pairs (*33, 34*); or poly-ADP-ribosyltransferases Tankyrase (TNKS), ANKS3, and DAG kinase SAM domains, which form left-handed helical filaments through extended ML-EH interactions (*23, 24, 35, 36*). Using the SSM server to compare structures showed that the ASK3 SAM domain aligns well (RMSD 1.4–2) with a range of the aforementioned SAM domains (Supp. Fig. 4). In order to understand the different oligomerisation propensity within ASK1–3 SAM domains, we compared sequence conservation and electrostatic potential of their ML and EH surfaces. A clear pattern emerges when modelling the ASK1 and ASK2 SAM domains based on ASK3 and mapping surface electrostatics (Figure 5A). The ASK3 ML surface is strongly negatively charged and EH surface is strongly positively charged, generating a highly complementary electrostatic interaction. In contrast, the ML surface of ASK2 has a generally hydrophobic character. Paired with a mildly positive EH surface it becomes apparent that the ASK2 ML and EH surfaces are not particularly compatible, hence the ASK2 SAM domain is generally monomeric. Instead, the ASK2 ML surface appears more complementary to the EH surface of the ASK1 SAM domain (Figure 5A), and mildly positive EH surface of ASK1 complementary to the mildly positive charge of the ASK1 ML surface. Such character would explain the observed behaviour of these domains in solution: where homotypic ASK1, or homotypic ASK2, interactions are transient and limited, whereas heterotypic interactions between ASK1/ASK2 surfaces are more complementary, readily leading to stable oligomer formation.

**Figure 5:**
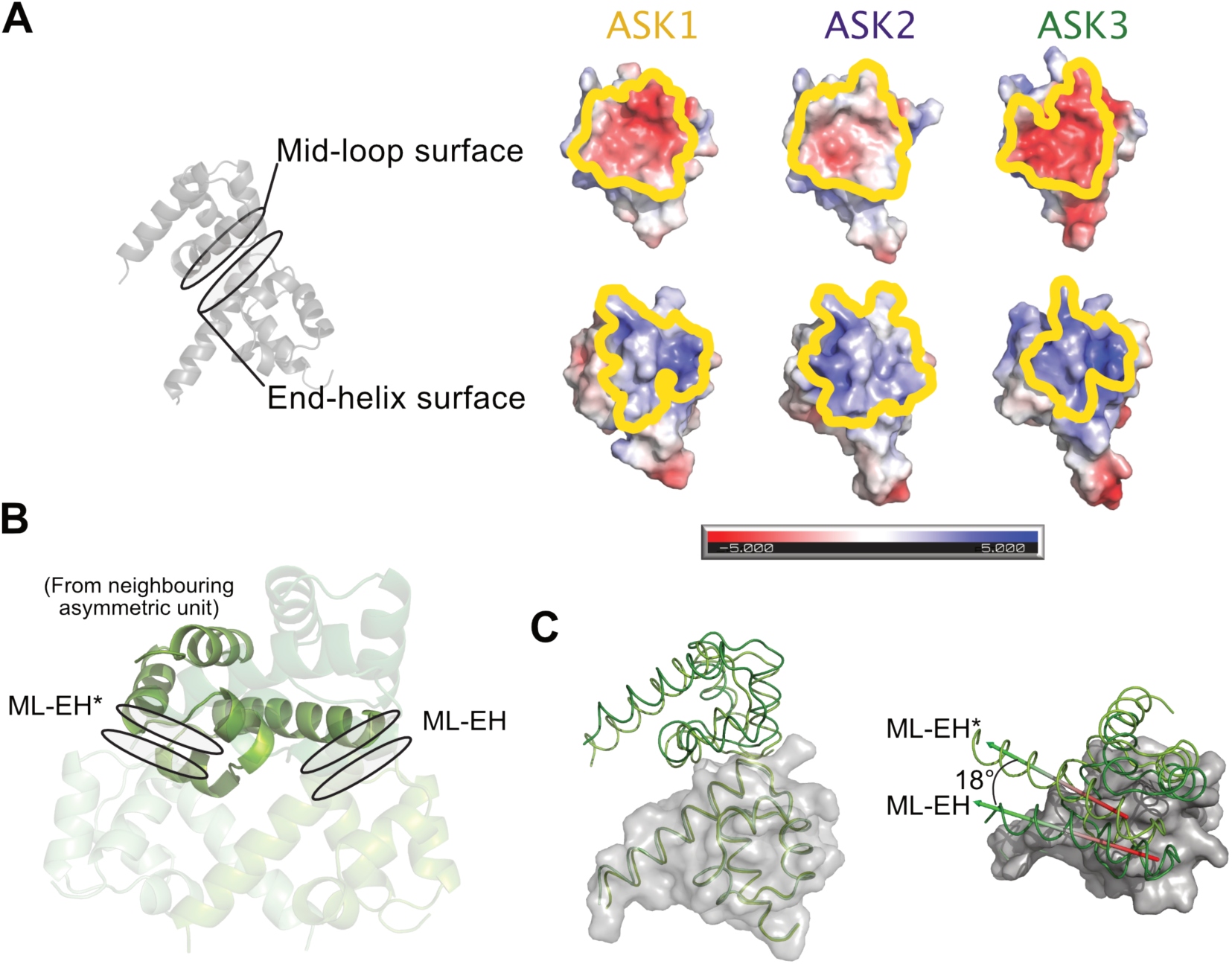
The ASK SAM domain ML-EH interface **(A)** Left: Schematic view of the ML-EH interface. Right: Electrostatic surfaces of ASK1–3 SAM domains (as calculated using APBS (37), with regions predicted to participate in ML-EH contacts outlined in yellow. Models of ASK1 and ASK2 were generated using MODELLER, based on the ASK3 SAM domain solved here. **(B)** Illustration of the ML-EH interface seen within the ASK3 asymmetric unit (ML-EH) and the similar but slightly offset arrangement with a crystallographically related SAM domain (ML-EH*) **(C)** Comparison of the ML-EH and ML-EH* interfaces. Pairs of SAM domains participating in each type of interface overlaid based on the bottom SAM domain. The top SAM domain is offset, quantitated by an 18° shift of the α5 helix.

While experiments in cells suggest that the ML-EH surface is crucial to ASK SAM domain function, and surface comparisons provide a basis for selective oligomerisation by ASK SAM domains, an outstanding question remains—why do ASK SAM domains form distinct soluble oligomers, rather than a continuous filamentous structure observed for other SAM domains? For example, mixing high-concentrations of purified monomer/dimer ASK1 and ASK2 SAM domains forms a distinct (non-filamentous) oligomer observed by various measures (Figure 3), and the ASK3 SAM domain also has a defined oligomeric state in solution. In considering this question we further analysed the crystallographic contacts in the ASK3 SAM structure, and observed a second, slightly offset ML-EH interface formed with a SAM domain from a neighbouring asymmetric unit (which we term ML-EH*; (Figure 5B)). The ML-EH* interaction uses effectively identical residues to ML-EH, but is offset by approximately 18° when considering the position of the α5 helix (Figure 5C). This indicates that there is malleability at the interface that could affect behaviour in solution, in line with other ML-EH complexes and filamentous assemblies.

### Solution conformation of ASK SAM oligomers

To investigate why the ASK SAM domains form higher-order oligomers of defined size we turned to small-angle X-ray scattering. We mainly sought to determine whether the SAM domains form an extended helix, or a more compressed helix or ring that may self-limit and give rise to a defined oligomer. In order to estimate experimental and actual scattering of oligomers formed through the ML-EH interface we considered three basic scenarios: oligomers formed by purely the ML-EH interface, purely the ML-EH* interface, or a mixture of the two. Modelling complexes formed through either ML-EH and ML-EH* have markedly different dimensions (Figure 6A), amplifying modest differences in the pairwise interaction (Figure 5C). Namely, the pure ML-EH oligomer forms an extended helix with a pitch of 52 Å and 7 units per turn, the purely ML-EH* interface a near-symmetrical closed ring, and the mixed interface an intermediate between the two helices (Figure 6A).

**Figure 6:**
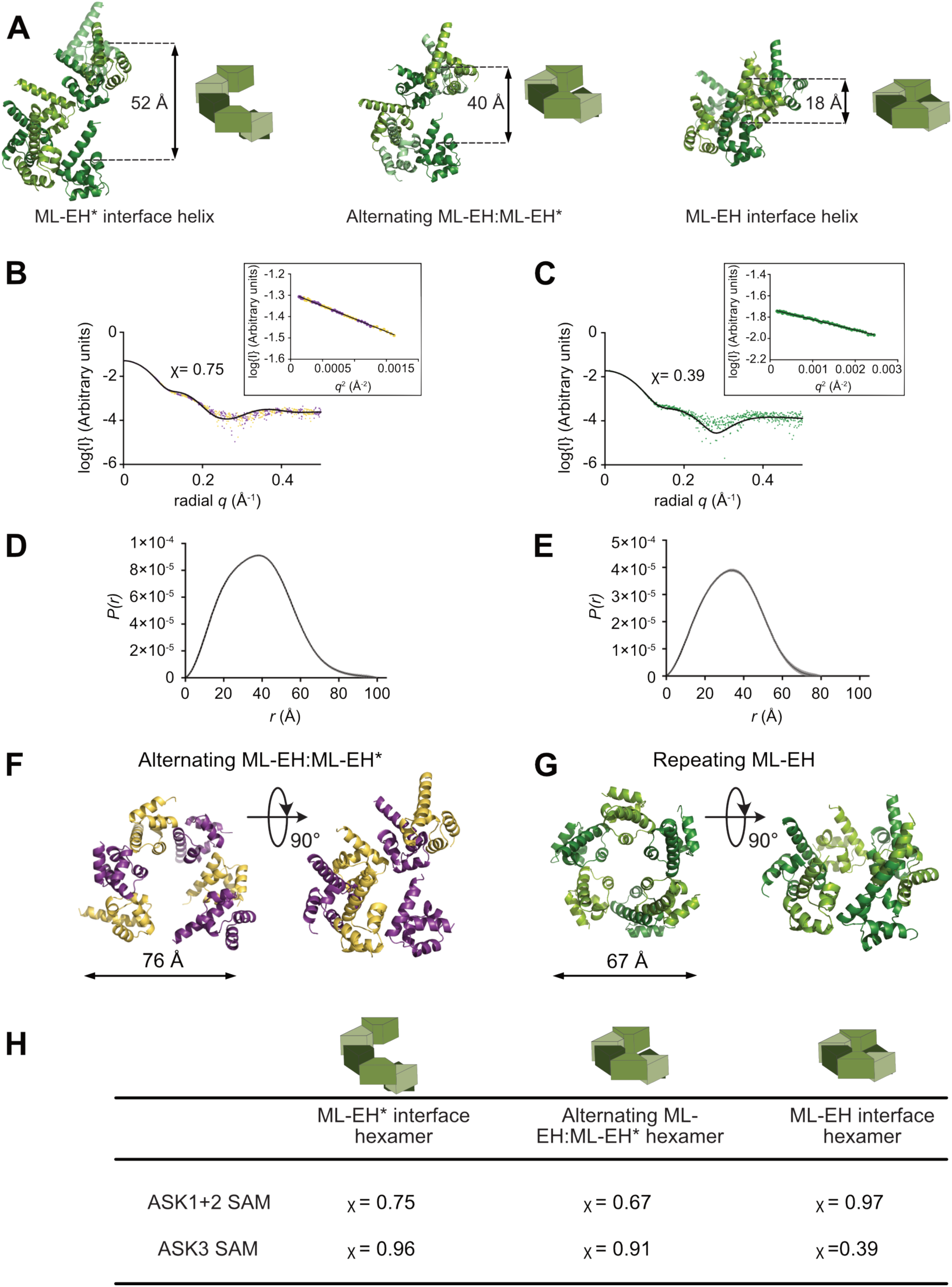
SAXS analysis of ASK1+2 and ASK3 SAM domains. (A)Schematic illustrating the helicies formed from the different ML-EH interfaces seen in the crystal lattice. (B/C) Experimental scattering curves with the best CRYSOL modelled fit (black line) and the Gunier plot (inset) for, ASK1+2 SAM (B); and ASK3 SAM (C). (C/D) Distance distribution plots for, ASK1+2 SAM (D); and ASK3 SAM (E). (F/G) Side and top views of best fit models for the (F) ASK1+2 SAM hexamer (alternating ML-EH/ML-EH*) and (G) ASK3 SAM hexamer (repeated ML-EH; within asymmetric unit). (H)Summary table of the fit of each model to the experimental SAXS data.

We collected small angle X-ray scattering data for various relevant ASK SAM domains: the isolated ASK1 and ASK2 SAM domains, the ASK1-ASK2 oligomer, and the stable ASK3 SAM domain oligomer (Table 2). Consistent with other in-solution data, the ASK2 SAM scattering data clearly fit a monomeric model (χ = 0.60; Supp. Fig. 5; Supp. Table 3). The experimentally determined scattering profile for ASK1 SAM domain was evaluated against both monomeric and dimeric models, whereby the radius of gyration determined experimentally from the *P(r)* plot (*P(r)R*_*g*_ *=* 14.3) is most consistent with that of the ML-EH dimer (Supp. Fig. 5;Supp. Table 3). A *P(r)R*_*g*_ of 14.3 is intermediate between the theoretical values for the monomer (*R*_*g*_ *=* 13.6) and MLEH dimer (*R*_*g*_ *=* 16.5), consistent with a mixed population observed in AUC and size-exclusion chromatography (Figure 1; Supp. Table 1).

High quality scattering data was collected for both the ASK1-ASK2 and ASK3 SAM oligomers, with the low *q* regions of the Guinier plots indicating homogenous, monodisperse protein samples (Figure 6C/F). Several conformational arrangements of pentamer, hexamer and heptamer were tested against experimentally measured scattering using CRYSOL (Supp. Fig. 5; Supp. Table 3; (*38*)). For all larger complexes the best fits to scattering data were clearly hexameric—clarifying ambiguous estimates of molecular weight arising from SEC-MALS and AUC (Figure 6). When considering flexibility of the ML-EH interface illustrated in Figure 6A, the best fits for ASK1-ASK2 and ASK3 differed. The best fit for the ASK1-ASK2 complex was an intermediate helix formed by a mixture of ML-EH and ML-EH* interfaces (χ =0.67), whereas ASK3 SAM clearly fit the most compact model tested (χ =0.39; Figure 6H). A compact, near-closed ring, for ASK3 is consistent with *D*_*max*_ estimates for ASK3 (80 Å), relative to 100 Å for ASK1-ASK2 (Figure 6; Table 2), and provides a clear mechanism for self-limiting oligomerisation. While slightly more extended than ASK3, filament formation by the ASK1-ASK2 complex is likely to be prevented through steric hindrance between neighbouring SAM domains.

## Discussion

MAP kinase signaling cascades are used throughout eukaryotes to translate external stimuli into cellular responses. Having a three-tiered phosphorylation cascade allows for both signal amplification, and for various levels of regulation. While MAP2K and MAPKs are relatively well conserved, MAP3Ks are significantly more divergent in their domain structure, made necessary by the diverse signals that MAP3Ks sense and respond to—from proliferative signals to signals eliciting cell death. One key mechanism of MAPK regulation is scaffolding of higher order complexes, which tethers relevant proteins into coherent signaling packages (*39*). Protein scaffolding can also modulate catalytic activity of kinases within MAPK pathways (*40, 41*). ASK kinases in humans are a three-membered sub-family of MAP3Ks, which have long been known to form higher order complexes that are inherent to their function. Here we demonstrate that ASK1–3 contain a previously uncharacterised domain at their extreme C-terminus, in the form of a SAM domain—a prevalent protein-protein interaction domain used throughout Eukaryota (*42*). The SAM domains from the three ASK orthologs have relatively divergent oligomerisation tendencies, even though they use the same oligomerisation surface as each other and many other SAM domains. The preferred state of ASK1–3 SAM domains varies, but notably does not extend beyond a hexameric state by any of the measures we tested, even at very high protein concentration. Both the formation of discrete oligomers, and preferential hetero-oligomer formation are notable features of ASK SAM domain complex formation.

SAM domains have been characterised as either monomeric or oligomeric (*43*), with oligomers generally exhibiting either pairwise dimer formation, or filamentous oligomer formation via the ML-EH interface. The relative orientation between ASK3 SAM domains in our crystal structure are comparable to that seen in either discrete or filamentous SAM domain oligomers. This translates to a roughly equivalent putative helical pitch (33–53 Å) to that of classical filamentous SAM domains, such as that of Tankyrase, Diacylglycerol kinase and others (Supp. Fig. 6;(*23, 24, 35, 44, 45*)). However, within the crystal there is obvious flexibility at the ML-EH interface, which is in line with SAXS analysis demonstrating relatively more-, or less-, extended quasi-helical structures formed by discrete ASK3 homo-hexamers, or ASK1-ASK2 hetero-hexamers. Nonetheless, it remains unclear why the ASK SAM domain oligomers are self-limiting and discrete, even at concentrations exceeding 300 µM. While a closed hexameric-ring as a mechanism of self-limitation is a tempting proposition, strong evidence is still lacking. For instance, a near-closed ring is the best match for experimental SAXS data from ASK3 SAM, but such a closed hexamer is not seen within the crystal structure. As the data stand, we hypothesise that either a closed ring, or steric occlusion—through flexibility at the ML-EH interface or though the variable C-terminal tails of each SAM domain—cause the ASK SAM domains to form discrete, self-limiting oligomers. Another clear point for future investigation is how other structural elements found in full-length ASK proteins, such as the kinase domain (that itself has been shown to dimerise) and the predicted C-terminal coiled coil (directly N-terminal to the SAM), might influence the oligomerisation behaviour of ASK1–3. Pertinent to this, the Kinase Suppressor of Ras (KSR) also contains a coiled-coil SAM domain arrangement. However, the CC-SAM of KSR is responsible for membrane association and/or scaffolding interactions with RAF MAP3Ks (*46, 47*), rather than higher-order oligomerisation. Nonetheless, our experiments in cells clearly show that the SAM domains play a major role in setting the stoichiometry of ASK signalosomes owing to ML-EH mutants exhibiting diminished stress-stimulated signalling. Previous studies of ASK1 incidentally bearing mutations (deletion or alanine mutations of Glycine1356/1357) of a similar surface also showed abrogated signalling in response to hydrogen peroxide, supporting a crucial role of the ML-EH surface in ASK signalling (*27*).

With growing knowledge of the domain structures of ASK kinases the obvious challenge is understanding how oligomerisation by the SAM at the C-terminus integrates with the raft of other interactions through their N-termini. Previously, we reported the crystal structure of the central regulatory region of ASK1—found N-terminal to the kinase domain—which links the N-terminal thioredoxin-binding domain to the kinase domain (*18*). A pleckstrin-homology domain within this novel fold appears to promote substrate MAP2K phosphorylation, which could occur on an intra- or inter-molecular basis. Other partners also have distinct oligomeric states. For instance, Peroxiredoxin-1 has been demonstrated to transduce peroxide signals to ASK1 (*48*), and peroxiredoxin proteins frequently adopt ring-shaped decamers/dodecamer of five or six Prdx-1 dimers (*49*). Similarly, the phosphatase PGAM5 is known to target ASK1 (*50*), and structural studies have shown that PGAM5 forms dodecameric rings that are important for its activity on an ASK1 substrate peptide (*51, 52*). N-terminal regions of ASK1 have been shown to interact with TNF receptor-associated factor type ubiquitin ligases (*17*), which form oligomers (*53*), and the F-box Cullin-ubiquitin ligase component Fbxo21 (*31*). Interestingly, the F-box protein FbxW11/βTrCP2 was also identified in our BiCAP analysis (Supp. Table 2), further reinforcing cross-regulation between ASK complexes and the ubiquitin proteasome system (*54*). Fbxo21 promotes Lys29-linked ubiquitination on ASK1 during viral infection (*31*), on lysine residues near the binding site for 14-3-3 proteins. 14-3-3 proteins are themselves dimeric regulators of ASK proteins that bind C-terminal to the kinase domain (*19*). Ultimately, many of the regulatory interactions of ASK kinases could be exacerbated, or compete, in the context of full-length proteins that are tethered through their C-termini. Thus, there are multiple mechanisms by which ASK regulation—either autoinhibition or transactivation—stand to be enhanced by SAM domain-based oligomerisation.

Preferential hetero-oligomerisation between the ASK1 and ASK2 SAM domains is a relatively simple molecular mechanism to explain the greater efficacy in eliciting stress responses in various settings (*4–8*). With isolated SAM domains the heterocomplex appears to be more stable at equivalent concentrations, which if translated to full-length proteins would mean that higher-order active complexes form more readily and are more persistent. Several other additional questions remain; including whether other SAM domain containing proteins may also be able to participate in ASK SAM oligomers. There was some incorporation of ASK3 into ASK1 complexes isolated during BiCAP, however, no other obvious SAM domain-containing candidates were identified (Supp. Table 2). Addressing the propensity of the isolated ASK3 SAM domain to bind ASK1/2 SAM domains; GST pulldowns suggest that the monomeric ASK2 SAM domain could be bound by GST-ASK3 SAM domain, but the reciprocal interaction occurs less readily (Figure 3A). Such behaviour might be explained by the stability of the ASK3 complex over a large concentration range. In equivalent experiments ASK1–ASK3 interactions appear less likely. However, whether ASK3 actively participates in endogenous ASK1/2 complexes in cells is a relevant functional question.

Overall, this study uncovers a common protein-protein interaction domain that plays an important role in the function of ASK kinases—adding to the conserved repertoire of functional domains found in MAP kinases, and their scaffolding proteins, from yeast through to humans. These findings reinforce modularity of signalling cascades in eukaryotes, in a manner that maintains remarkable specificity despite structural similarity. ASK kinases appear to be particularly rich in autoregulatory interaction domains, in line with their role at the intersection of many cellular stresses. Understanding how the features work in concert, at the protein level and in cells, remains an ongoing challenge relevant to a multipurpose signalling hub.

## Acknowledgements

We thank members of the Mace Laboratory for useful discussions and assistance, particularly Pavel Filipcík for X-ray crystallography assistance. Various plasmids were kind gifts from Daniel Liebler, William Hahn, David Root, and Stephen Michnick (See Methods). This work was funded by the Marsden Fund of New Zealand; PDM was also supported by a Rutherford Discovery Fellowship administered by the Royal Society of New Zealand. Support is also acknowledged from the Victorian State Government Operational Infrastructure Support, NHMRC IRIISS grant (9000433), and an NHMRC fellowship (1105754; to J.M.M.). This research was undertaken in part using the small-angle X-ray scattering and the MX2 beamline at the Australian Synchrotron, part of ANSTO, and made use of the ACRF detector. We thank the New Zealand synchrotron group for facilitating access to the MX beamlines, and the Biomolecular Interaction Centre for AUC access.

## Materials and Methods

### DNA Constructs

Tandem-tagged constructs for HEK293 expression and HNE induction (ASK1 (HA-Flag) and ASK2 (HA-V5) Addgene # 69726, #69727, respectively) were a kind gift from Daniel Liebler (*28*). For BiCAP experiments The pDONR223-MAP3K5 used (Addgene plasmid 23517) (*55*)) was a kind gift of Dr William Hahn and Dr David Root. An expression vector encoding full length Venus fluorescent protein was a kind gift from Dr Stephen Michnick (University of Montreal). The ASK1 SAM domain was amplified from the MegaMan Transcriptome library (Agilent). Constructs comprising ASK2 and ASK3 were amplified from Addgene plasmids (#69727 and #69728, respectively). Indicated fragments were cloned into a pET-LIC vector either containing an N-terminal 6xHis or GST tag, and a 3C protease cleavage site. All mutants were generated using quikchange mutagenesis and verified by Sanger sequencing.

### Protein Expression and Purification

All recombinant proteins were expressed in *E. coli* BL21(DE3) in LB media, induced with IPTG overnight at 18 °C, and lysed by sonication. ASK1 (1290–1374), ASK2 (1216–1288) and ASK3 (1241–1313) were initially purified from clarified *E. coli* lysate by Ni^2+^-affinity chromatography using HisSelect resin (Sigma), followed by size exclusion chromatography (Superdex 75 column; GE Healthcare), with a 3C cleavage step between. SEC was carried out using a buffer consisting of 10 mM HEPES (pH 7.6), 150 mM NaCl and 2 mM DTT. Purified proteins were snap frozen in aliquots using liquid nitrogen.

### Analytical Ultracentrifugation

Sedimentation velocity experiments using absorbance optics were conducted in a Beckman XL-I analytical ultracentrifuge. Initial scans were performed at 3,000 rpm to determine the optimal wavelength for data collection. Experiments were conducted at 20 °C, the pre-determined wavelength, continuous mode, 50,000 rpm in 20 mM HEPES (pH 7.6), 150 mM NaCl 0.2 mM TCEP. Buffer density and viscosity and an estimate of the partial specific volume of proteins (v-bar) was calculated using SEDNTERP. Data were fitted to a continuous sedimentation coefficient [c(s)] model using SEDFIT. Data were visualised by creating c(s) vs. s graphs using the GUSSI software.

### Crystallisation and Structure Solution

ASK3(1241–1313) was initially crystallised in 0.1 M Bis-Tris pH 6.5, 25 % (w/v) PEG3350 at a 1:1 drop ratio. Optimisation was carried out using the Hampton Research stock options pH kit with diffracting crystals grown in 0.1 M sodium citrate tribasic dihydrate pH 5, 25 % (w/v) PEG3350, and frozen with the addition of 20% (v/v) glycerol. X-ray diffraction data were collected at the Australian Synchrotron beamline MX2. Native and iodide soaked (0.5 M NaI) crystals were collected at 0.9357 and 1.456 Å wavelengths, respectively. The structure was solved using single-wavelength anomalous dispersion, using a 1.8 Å dataset. The Auto-Rickshaw webserver was used for structure solution by generating initial phases and an electron density map (*56*). An initial model was built by Buccaneer and improved using cycles of automated and manual refinement using the PDB_REDO web server (*57*), Phenix (*58*) and Coot (*59*). Structural figures were generated using PyMOL (Schrodinger).

### SEC-MALS

SEC-MALS was conducted using a Wyatt Dawn 8+ detector (Wyatt Technology) coupled to a Superdex 75 10/300 column (GE Healthcare) and a refractive index detector. Samples were run in 10 mM HEPES (pH 7.6), 150 mM NaCl, and 0.3 mM TCEP and loaded at 100 or 200 uM. All data were analyzed using ASTRA V software.

### Cell Lines and Cell Culture

HEK-293FT cells were grown in DMEM (Life Technologies, 10566) supplemented with 10% fetal bovine serum (Sigma-Aldrich, F8067), 2 mM L-glutamine (Life Technologies, 25030081), 100 units/mL Penicillin-Streptomycin (Life Technologies, 15140122), Non-Essential Amino Acids (Hyclone, SH30238) and 1 mM Sodium Pyruvate ((Hyclone, SH3023901). Cells were grown at 37 °C in a humidified atmosphere with 5% CO_2_.

### HNE Stimulation

Cells were transiently co-transfected with Lipofectamine 3000 (Life Technologies, L3000015). A total of 3 μg plasmid DNA was used; either 1.5 μg of relevant ASK construct and/or pcDNA3. Cells were grown for twenty-four hours before 4-hydroxy-2-nonenal (HNE) treatment. One hour before treatment the medium was replaced with serum free-media. Cells were treated with either ethanol (vehicle control) or 50 µM HNE for 1 h at 37 °C. Cells were harvested into the treatment medium and centrifuged at 500 × *g* for 5 min at 4 °C. The cell pellets were washed twice with ice cold phosphate buffered saline (PBS) and the cell pellet resuspended in 100 μL of 4× SDS-PAGE Sample Buffer. Samples were frozen in liquid nitrogen and stored at −80 °C until use.

### Western Blotting

For analysis by western blot, samples were separated by SDS/PAGE and transferred to 0.2 µm nitrocellulose (Life Technologies, IB23002). Membranes were blocked in 5% BSA (w/v) in TBS-T. Membranes were incubated with primary antibodies overnight at 4°C in 5% BSA (w/v) in TBS-T. Antibodies used in this study were rabbit monoclonal p38 MAPK (1:2000, CST, #8690), rabbit monoclonal Phospho-p38 MAPK (Thr180/Tyr182) (1:500, CST, #4511), Phospho-ASK1 (Thr845) (1:1000, CST, #3765), mouse monoclonal V5 tag (1:5000, Abcam, ab27671), rabbit monoclonal DYKDDDDK tag (Flag tag, 1:1000, CST, #14793) and/or mouse monoclonal α-tubulin (1:10,000, Millipore, 05-829). Following three washes with TBS-T, membranes were incubated with secondary antibodies diluted in TBS-T with 1% (w/v) BSA for 1 hour at room temperature. Secondary antibodies used were goat anti-rabbit IRdye 680LT (LI-COR), goat anti-mouse IRdye 800LT (LI-COR) or goat anti-rabbit HRP-conjugated (1:10,000, Abcam, ab6721; used with Phospho-p38 MAPK). Membranes were washed a further three times with TBS-T. Membranes were then developed with the Odyssey Fc imaging system.

### Bimolecular complementation affinity purification

Vectors expressing V1 or V2 tagged fusions of ASK1 were generated by recombination cloning into pDEST-V1 or pDEST-V2 destination vectors using Gateway LR Clonase enzyme mix (Life Technologies) according to manufacturer’s instructions and verified by sequencing.

HEK293T cells were grown in 10 cm dishes and transfected with 2.5 μg of each BiCAP construct using JETprime transfection reagent (Polyplus). After 16 h, cells were harvested by washing twice with warm PBS and then scraping on ice with ice-cold lysis buffer (50 mM Tris HCl pH 7.4, 150 mM NaCl, 1mM EDTA, 1% (v/v) Triton X-100) supplemented with fresh EDTA-free protease inhibitor cocktail and 0.2mM sodium orthovanadate. Samples were cleared by centrifugation at 18000g for 10 min at 4°C to remove cellular debris, prior to proceeding with affinity purification using GFP-Trap_A agarose beads (ChromoTek GmbH), trypsin digestion and nanoLC-MS/MS as previously described in detail (*29, 30*).

### Small-Angle X-ray Scattering

SAXS data was collected using the SAXS/WAXS beamline at the Australian Synchrotron with an inline gel-filtration chromatography setup (*60–62*). Protein samples (ASK1 SAM 50 µL at 9.2 mg.mL^-1^; ASK2 SAM 90 µL at 8.5 mg.mL^-1^; ASK1+2 SAM 90 µL at 5.2 mg.mL^-1^; ASK3 SAM 50 µL at 8.3 mg.mL^-1^) were injected onto a Superdex 75 Increase 5/150 column and eluted in 10 mM HEPES pH 7.5, 150 mM NaCl, 5% glycerol and 0.2 mM TCEP at a flow rate of 0.5 mL.min^-1^. Protein was eluted from the column into a 1 mm diameter quartz capillary orthogonally aligned to the X-ray beam. The coflow system, providing sheath flow, was used to achieve stable laminar flow through the capillary reducing radiation damage (*63*). Data was collected at 285 K using an X-ray beam of 1.03 Å in wavelength and 2 s exposure times. X-ray scattering was measured by a Pilatus 1M or 2M detector (Dectris). Primary data reduction and buffer subtraction was performed onsite at the Australian Synchrotron using scatterBrain software developed in-house (Stephen Maudie, Australian Synchrotron). Data analysis was performed using Primus (*64*), GNOM (*65*), and Crysol (*66*) from the ATSAS package (*67*).

---

**Supp. Fig. 1:**
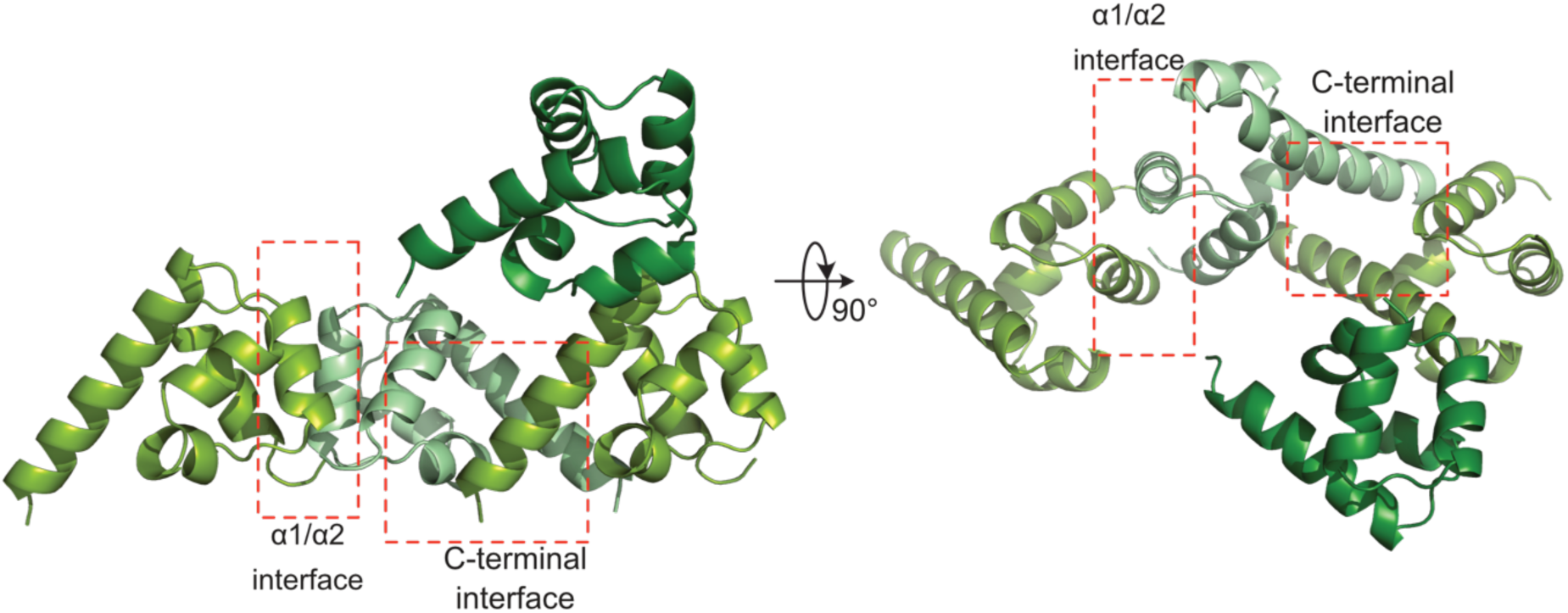
ASK3 SAM crystal packing. Cartoon Illustration of the crystallographic asymmetric unit, and one SAM domain from a neighbouring asymmetric unit that interacts at the α1/α2 interface. Interfaces referred to in the text are indicated

**Supp. Fig. 2:**
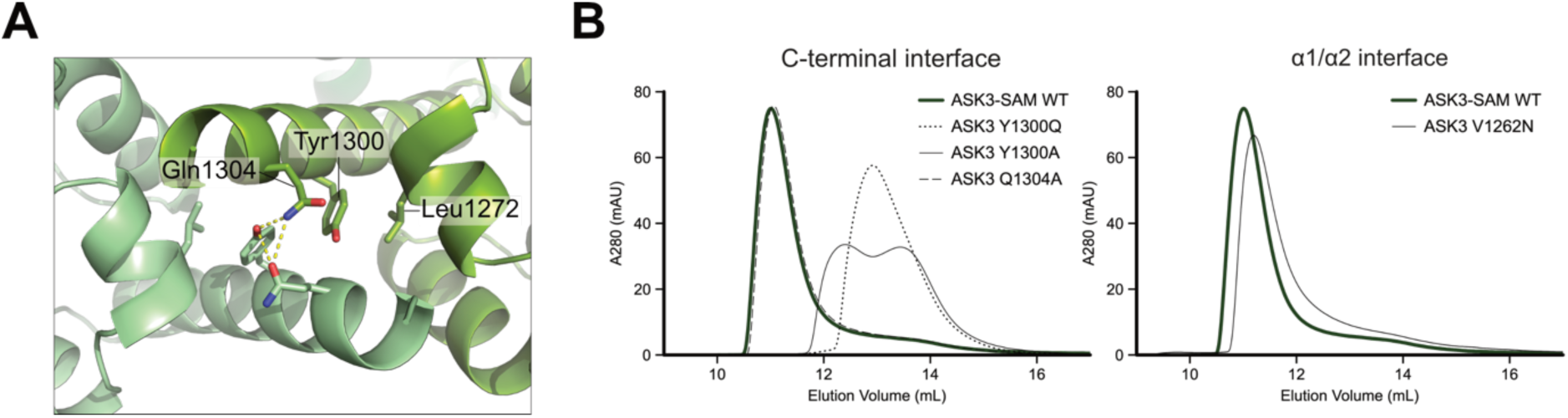
Analysis of ASK3 SAM mutants (A) closeup view of residues at the C-terminal interface. (B) Size-exclusion chromatography of the WT ASK3 SAM and mutations at the C-terminal (left) and α1/α2 (right) interface.

**Supp. Fig. 3:**
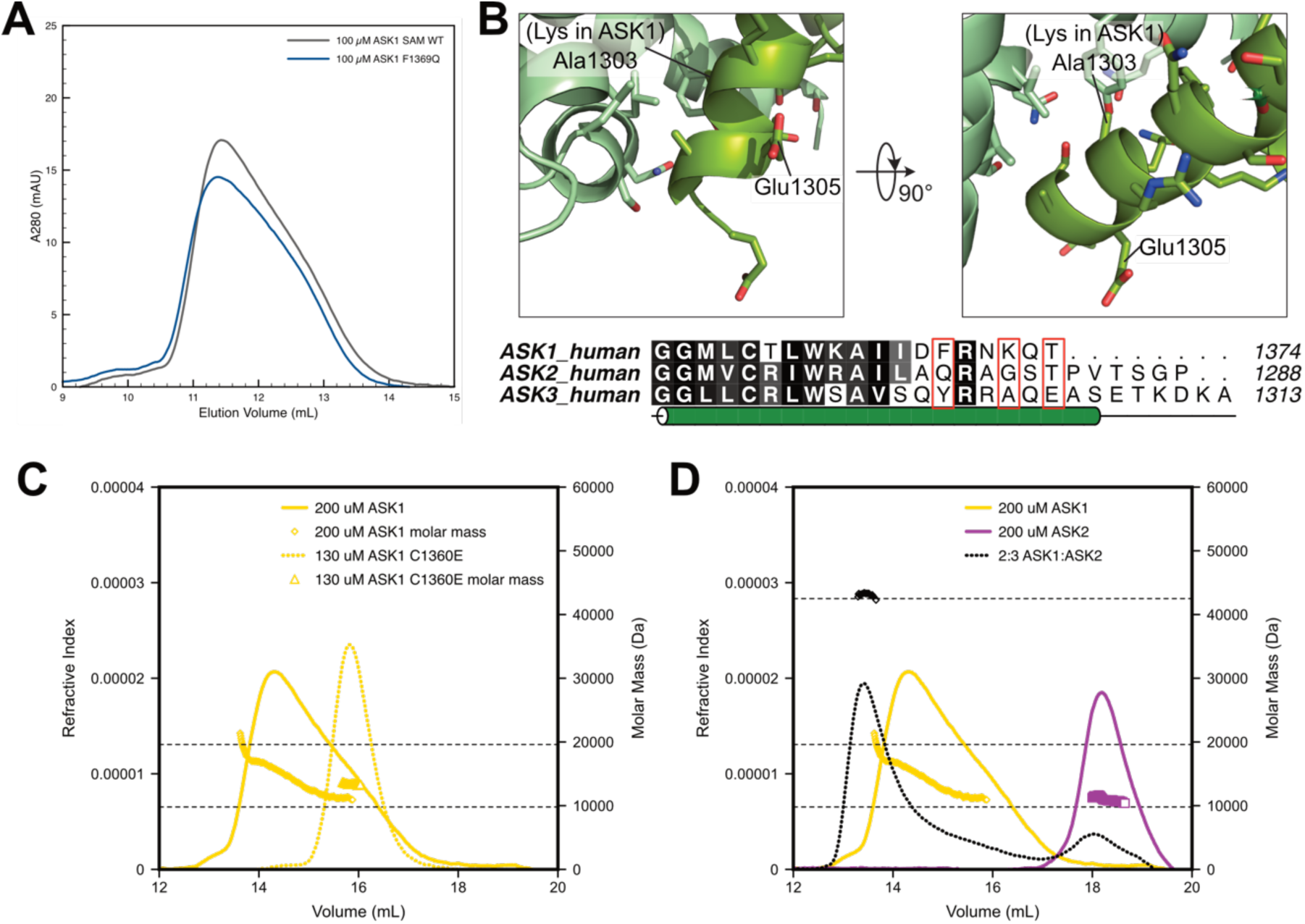
Potential ASK1 oligomer interfaces. (A) Size-exclusion chromatography of WT ASK1 SAM and F1369Q mutation (equivalent to ASK3 Y1300Q). (B) Closeup view of the ASK3 C-terminal tail interaction, relative to sequence conservation in ASK1. Indicated on the alignment are ASK1/ASK3 residues F1369/Y1300, Lys1372/Ala1303 (which would could not be accomodated in a putative ASK1 complex), and Thr1374/Glu1305, to indicate the the position of the final residue of ASK1. (C) SEC-MALLS of WT and C1360E mutation of ASK1. (D) SEC MALS of WT ASK1, ASK2 and a mixture of the two SAM domains, each at a total concentration of 200 µM.

**Supp. Fig. 4:**
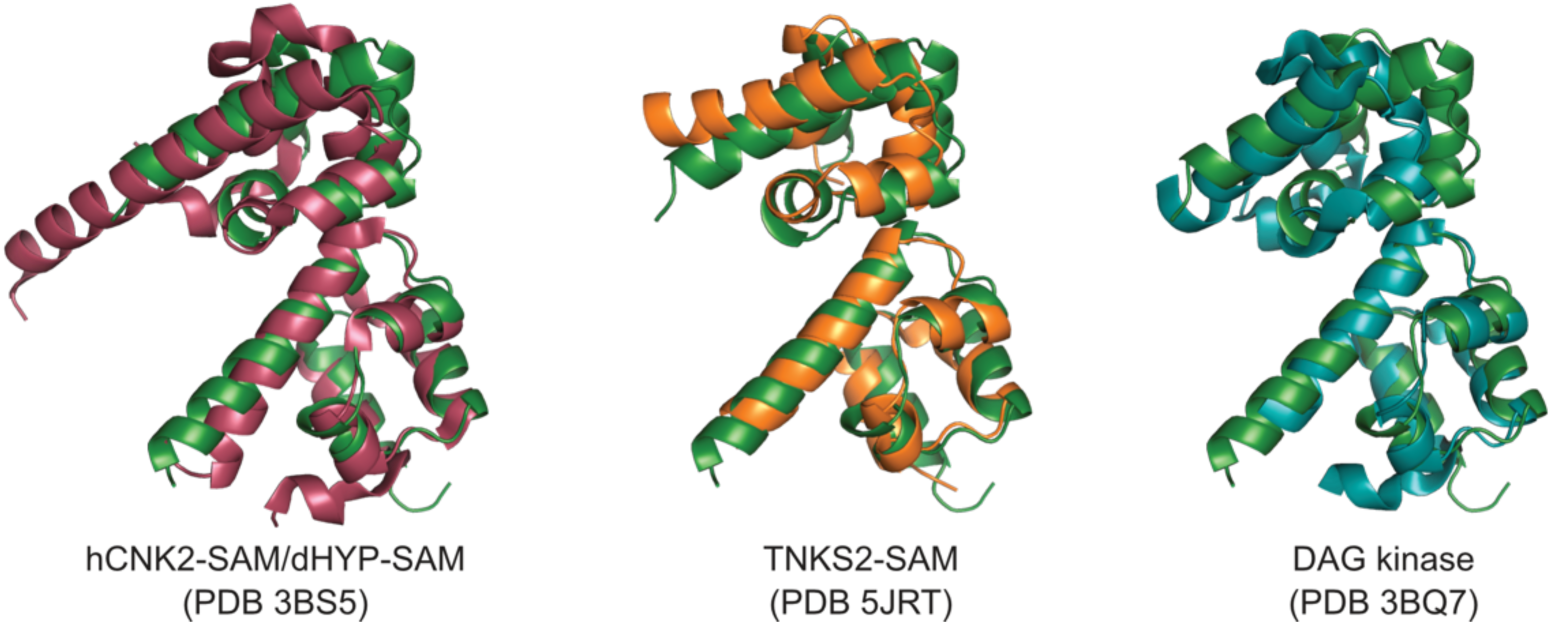
Comparisons of the ML-EH pairwise interaction of ASK3 with indicated SAM ML-EH structures.

**Supp. Fig. 5:**
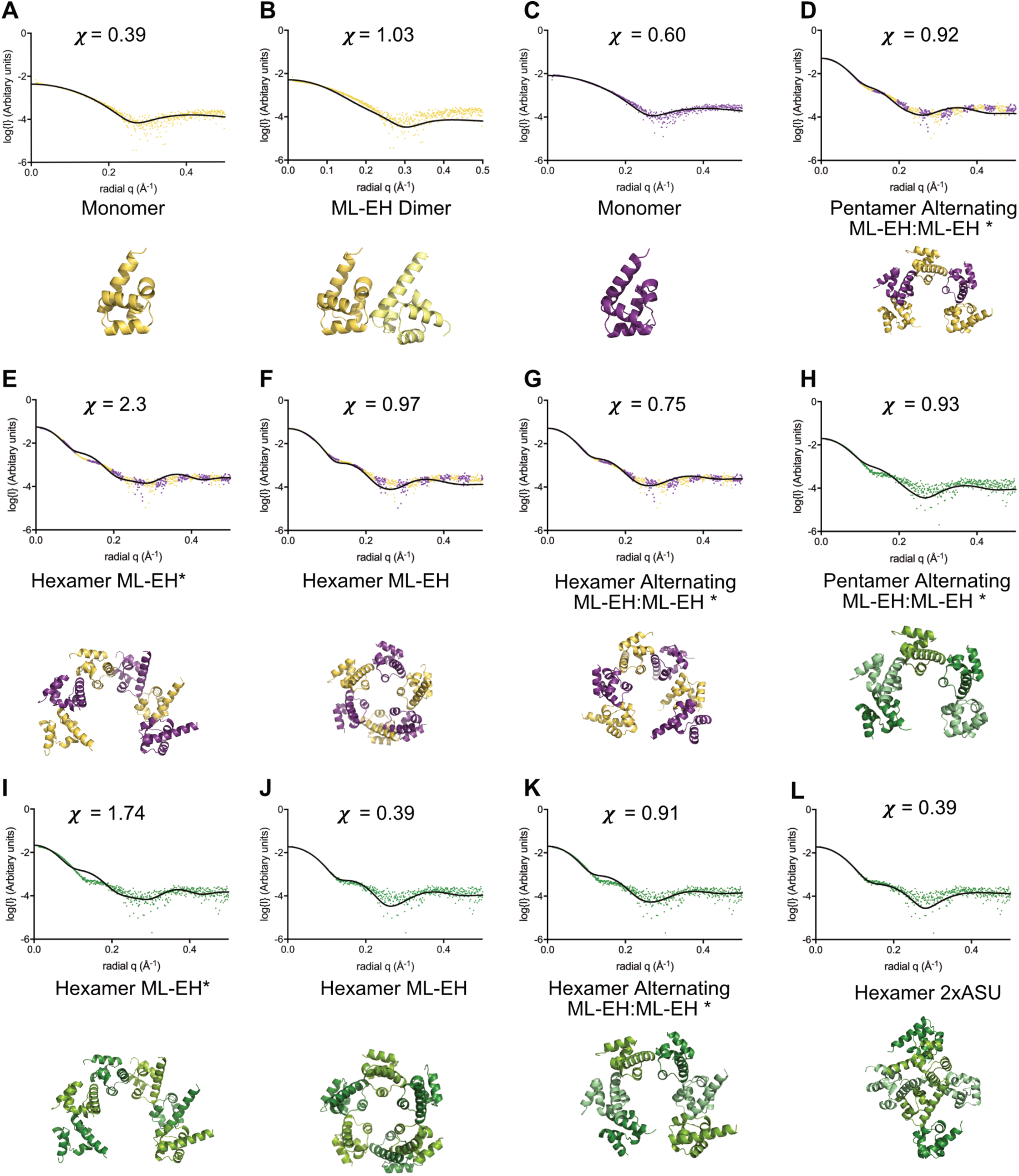
Comparison of SAXS models to scattering data. CRYSOL fits of models to experimental SAXS scattering data, with corresponding model illustrated below for (A/B) ASK1 SAM; (C) ASK2 SAM; (D–G) ASK1+2 SAM; (H–L) ASK3 SAM.

**Supp. Fig. 6:**
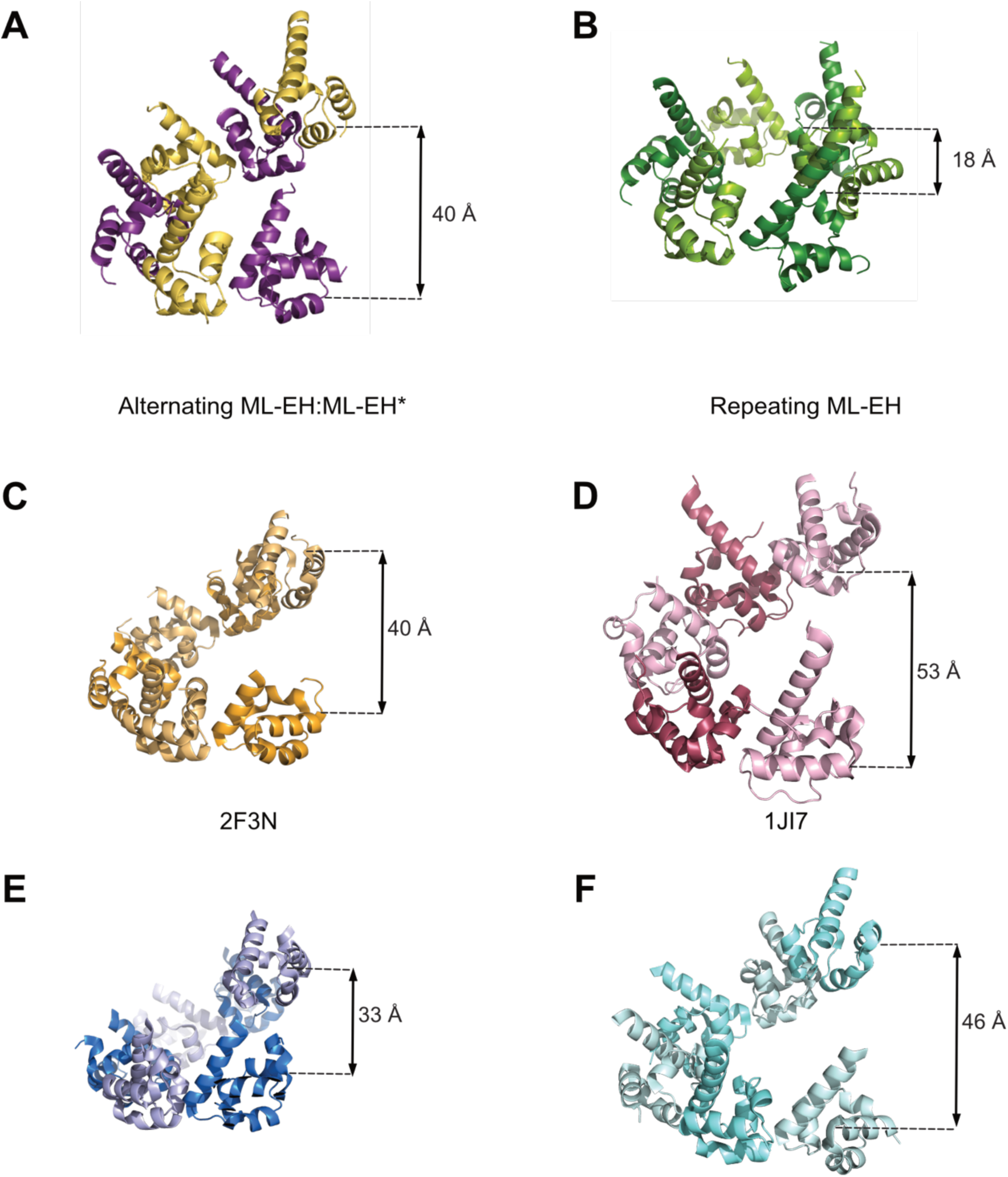
Comparison of the pitch of helical filaments formed by SAM domain ML-EH interfaces. (A) Putative ASK1-ASK2 alternating ML-EH:ML-EH*. (B) Putative ASK3 repeating ML-EH. (C) SHANK3 from Rattus norvegicus (PDB ID: 2F3N). (D) Human ETS-related protein TEL1 (PDB ID: 1JI7). (E) Human Diacylgylcerol kinase 1 (PDB ID: 3BQ7). (F) Human Tankyrase 2 (PDB ID: 5JRT).

**Supp. Table. 1:** Summary of AUC data. ^ Frictional ratio determined by SEDFIT. * Weight averaged sedimentation coefficients obtained from the c(s) distribution.

| | Concentration | | rmsd | $f/f_0$ ^ | Z score | Mass (kDa) | s* |
| --- | --- | --- | --- | --- | --- | --- | --- |
|  | mg/mL | μM |  |  |  |  |  |
| ASK1 SAM domain | 0.15 | 15 | 0.027 | 1.30 | 0.13 | 13.9, 35.2 | 2.05 |
|  | 0.3 | 31 | 0.028 | 1.50 | 1.19 | 14.5, 40.7 | 2.39 |
|  | 0.5 | 51 | 0.0093 | 1.27 | 0.02 | 33.7 | 2.77 |
|  | 1.5 | 153 | 0.010 | 1.42 | 1.34 | 41.9 | 2.85 |
| ASK2 SAM domain | 0.15 | 18 | 0.027 | 1.34 | 1.44 | 9.49 | 1.17 |
|  | 0.3 | 37 | 0.029 | 1.39 | 0.75 | 10.2 | 1.17 |
|  | 0.5 | 61 | 0.0097 | 1.46 | 0.03 | 11.7 | 1.26 |
|  | 1.5 | 183 | 0.010 | 1.35 | 3.96 | 10.6, 25.4 | 1.33 |
|  | 3 | 366 | 0.011 | 1.48 | 7.23 | 14.2, 31.4 | 1.64 |
| ASK3 SAM domain | 0.15 | 18 | 0.026 | 1.49 | 2.16 | 9.46, 47.0 | 2.76 |
|  | 0.3 | 36 | 0.027 | 1.42 | 0.42 | 9.36, 44.4 | 2.85 |
|  | 0.5 | 59 | 0.0097 | 1.35 | 1.44 | 41.3 | 2.92 |
|  | 1.5 | 177 | 0.010 | 1.40 | 1.02 | 44.3 | 2.95 |
| ASK1+ASK2 | 1.5+1.5 | 336 | 0.013 | 1.37 | 0.97 | 11.4, 47.5 | 2.81 |

**Supp. Table. 2:**
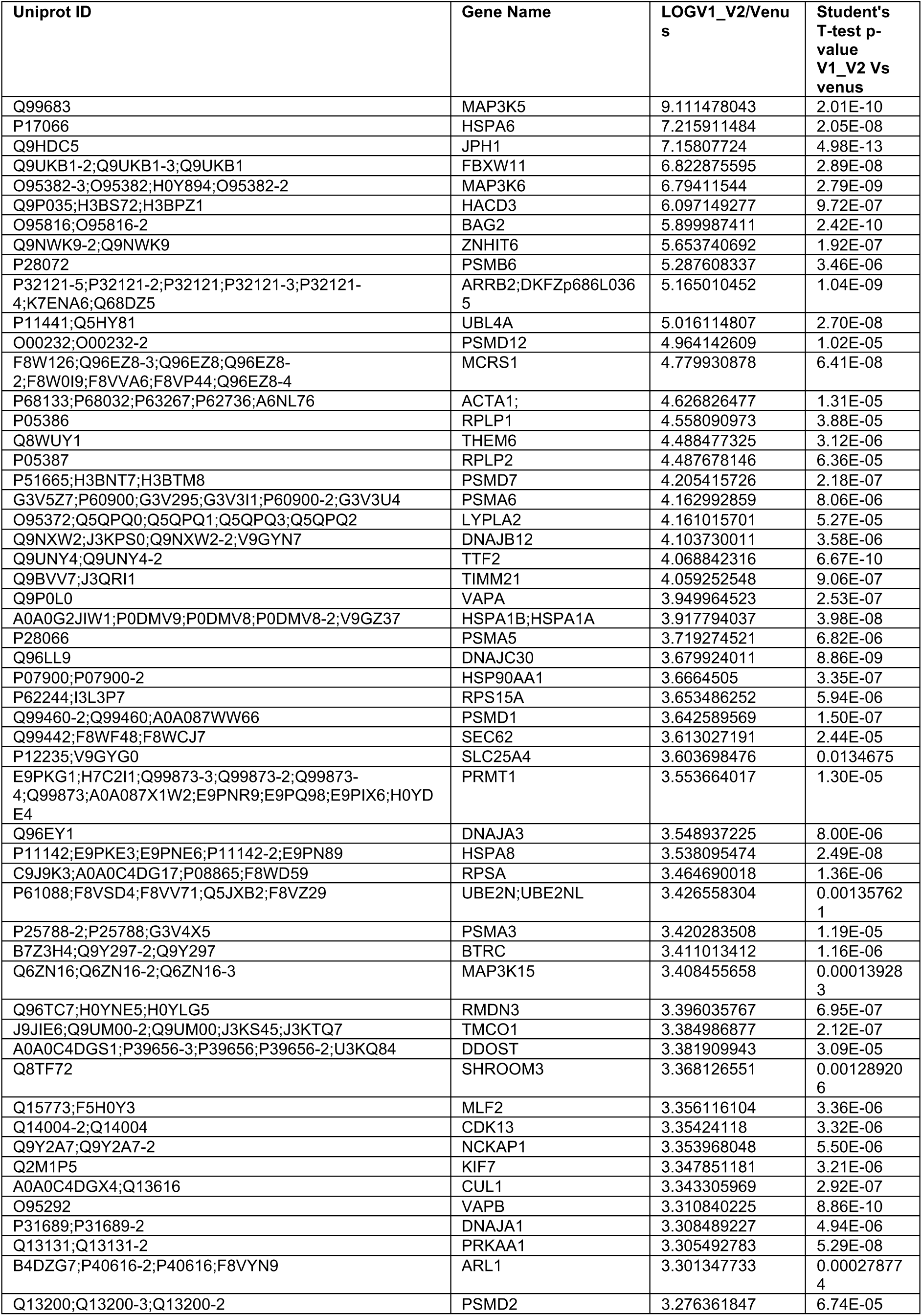
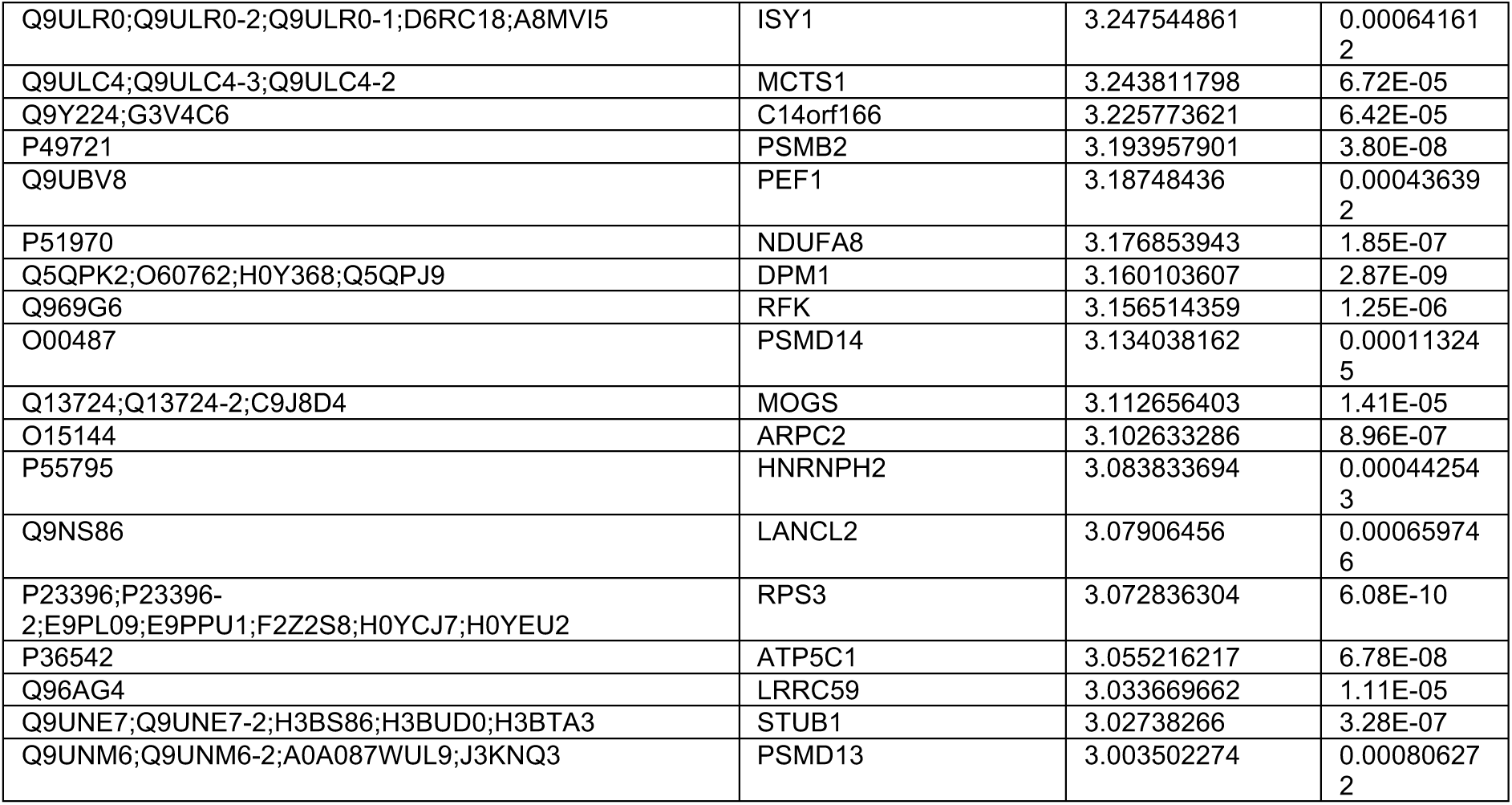
Summary of BiCAP mass spectrometry data

**Supp. Table. 3:** Summary of experimental SAXS data fit to possible oligomeric models

|  | Experimental Data |  | Models |  |  |  |
| --- | --- | --- | --- | --- | --- | --- |
| | $R_g$ | $D_{max}$ (Å) | Model | Theoretical $R_g$ | Diameter (Å) | Fit $\chi$ |
| ASK1 SAM | 14.3 | 40 | Monomer | 13.6 | 30 | 0.39 |
|  |  |  | ML-EH Dimer | 16.5 | 40 | 1.03 |
| | | | α3-4 Dimer | 18.5 | | 1.00 |
| | | | α5 Dimer | 17.2 | | 1.00 |
| ASK2 SAM | 16.0 | 65 | Monomer | 13.9 | 30 | 0.60 |
| ASK1+2 SAM | 29.2 | 100 | Alternating Pentamer | 29.0 | 78 | 0.91 |
|  |  |  | Alternating Hexamer | 28.6 | 77 | 0.67 |
|  |  |  | ML-EH Hexamer | 27.3 | 68 | 0.97 |
|  |  |  | ML-EH* Hexamer | 28.7 | 75 | 0.74 |
| ASK3 SAM | 25.8 | 80 | Alternating Pentamer | 27.6 | 78 | 0.93 |
|  |  |  | Alternating Hexamer | 27.4 | 77 | 0.91 |
|  |  |  | ML-EH Hexamer | 25.4 | 68 | 0.39 |
|  |  |  | ML-EH* Hexamer | 27.5 | 75 | 0.96 |

